# Higher stochasticity of microbiota composition in seedlings of domesticated wheat compared to wild wheat

**DOI:** 10.1101/685164

**Authors:** Ezgi Özkurt, M. Amine Hassani, Uğur Sesiz, Sven Künzel, Tal Dagan, Hakan Özkan, Eva H. Stukenbrock

**Affiliations:** Environmental Genomics, Christian-Albrechts University of Kiel, Am Botanischen Garten 1-9, 24118 Kiel; Environmental Genomics, Max Planck Institute for Evolutionary Biology, August-Thienemann-Str. 2, 24306 Plön, Germany; Department of Field Crops, Faculty of Agriculture, University of Çukurova, 01330 Adana, Turkey; Evolutionary Genetics, Max Planck Institute for Evolutionary Biology, August-Thienemann-Str. 2, 24306 Plön, Germany; Institute of General Microbiology, Christian-Albrechts University of Kiel, Kiel, Germany

**Keywords:** Microbial profiling, seedborne plant microbiota, endophytes, *Triticum aestivum*, domestication, co-adaptation

## Abstract

Plants constitute an ecological niche for microbial communities that colonize different plant tissues and explore the plant habitat for reproduction and dispersal. The association of microbiota and plant may be altered by ecological and evolutionary changes in the host population. Seedborne microbiota, expected to be largely vertically-transferred, have the potential to co-adapt with their host over generations. Reduced host diversity because of strong directional selection and polyploidization during plant domestication and cultivation may have impacted the assembly and transmission of seed-associated microbiota. Nonetheless, the effect of plant domestication on the diversity of their associated microbes is poorly understood. Here we show that microbial communities in domesticated wheat, *Triticum aestivum*, are less diverse but more inconsistent among individual plants compared to the wild wheat species, *T. dicoccoides*. We found that diversity of microbes in seeds overall is low, but comparable in different wheat species, independent of their genetic and geographic origin. However, the diversity of seedborne microbiota that colonize the roots and leaves of the young seedling is significantly reduced in domesticated wheat genotypes. Moreover, we observe a higher variability between replicates of *T. aestivum* suggesting a stronger effect of chance events in microbial colonization and assembly. We also propagated wild and domesticated wheat in two different soils and found that different factors govern the assembly of soil-derived and seedborne microbial communities. Overall, our results demonstrate that the role of stochastic processes in seedborne microbial community assembly is larger in domesticated wheat compared to the wild wheat. We suggest that the directional selection on the plant host and polyploidization events during domestication may have decreased the degree of wheat-microbiota interactions and consequently led to a decreased stable core microbiota.

## Introduction

Plants coexist with a large diversity of microorganisms. Most of the plant-associated microbiota is acquired from the environment, while a smaller component is vertically inherited e.g. via the seed (1), (2). Plant-microbe interactions range from parasitic or neutral to beneficial whereby the microbiota can contribute to increased nutrient uptake, stress tolerance and disease resistance (3), (4), (5). There is a growing attention on plant microbiota and its role in the future improvement of agricultural plant production. However, the underlying plant traits that govern plant microbial assembly and maintenance are still poorly understood.

Interactions and co-evolution of plants with their associated pathogens and mutualists have been intensively studied (6), (7), (8). It is well-known that the plant immune system plays a fundamental role in the interaction with both pathogens and mutualists (9), (10). Consequently, genes encoding immune related proteins co-evolve with microbial-produced proteins to either abort or facilitate interactions (11). To which extent plants co-evolve with their associated microbiota has so far little been addressed. Plant-microbe co-evolution may be pronounced for seed-associated microbes that co-exist with their host over multiple generations. Seeds constitute a microbial niche for dispersion and transmission over multiple host generations. Also, seedborne microbes may harbour competitive advantages compared to the environmentally introduced microbes because they are already established in the plant niche during early colonization of the plant. Knowledge on seed-associated microbes, notably seedborne fungi, is relatively limited compared to microbiota associated with other plant tissues such as leaves and roots (12), (13). This is partly due to technical challenges of handling single seeds and the extraction of sufficient amounts of microbial DNA. For example, the model species *Arabidopsis thaliana* produces very small seeds that has limited detailed studies of the seedborne microbiota in this species (14). Thus, most studies of seedborne microbial communities have used culture dependent techniques or pooled multiple seeds for culture independent methods (2), (13), (15), (16), (17). While these studies have provided insight into overall diversity of seedborne microbiota, they have not allowed high-resolution analyses of microbial diversity within individual seeds, nor to which extent these microbial taxa co-evolve with the plant host.

In this study, we have assessed microbial communities of individual seeds across several wild and domesticated wheat species. Specifically, we have asked to which extent the seedborne microbial communities of closely related plant species differ and investigated differences that may reflect divergent co-adaptation of the microbiota. We focused our study on wheat, which represents an ideal model system to study the impact of recent artificial plant selection associated with domestication and genetic plant divergence on plant associated microbiota. Bread wheat, *Triticum aestivum*, was domesticated in the Fertile Crescent 10-12.000 years ago and the domestication history has been well characterized (18), (19), (20). Moreover, the underlying genetics of wheat domestication has been described in details, including bottlenecks in the wheat diversity following strong directional selection and polyploidization (21), (20). More recently, comparative genome analyses have allowed identification of domestication signatures along the *T. aestivum* genome (22). *T. aestivum* has been dispersed worldwide with wheat cultivation and constitute a major crop on all continents (23). Wild relatives of the hexaploid wheat *T. aestivum* originate in the Near East and can be found in natural grassland vegetation, including tetraploid emmer wheat, *Triticum dicoccoides,* and diploid einkorn, *Triticum boeoticum* and red wild einkorn *Triticum urartu* (24), (25). The well-documented domestication history and close relatedness of wild and domesticated wheat provide an optimal framework for comparative analyses of plant associated microbial communities. Moreover, it allows us to address the consequences of plant domestication on seed-associated microbial communities.

We hypothesized that strong directional selection during wheat domestication has impacted genetic factors involved in microbial assembly, for example immune related genes. If the plant genotype exerts a considerable impact on the plant associated microbiota, we would then expect to observe differences in microbial diversity and community composition. To address this hypothesis, we focused our study on both bacterial and fungal endophytes of wheat seedlings. We show that domestication did not entail a loss of microbial diversity in seeds, but rather in the diversity of early colonizers of the domesticated plant. We also addressed the alteration of microbiota assembly when wheat seeds were propagated in soil and we demonstrate that soil is a main determinant of microbial diversity. Moreover, in leaves of wild wheat seedlings show less variation between replicates when grown in natural soil, consistent with the observation from axenic wheat seedlings. On the other hand, the soil type is a main determinant in microbial community composition. Finally, we also suggest that the biodiversity of wheat-associated fungal community is governed by different selection regimes in comparison to the bacterial community.

## Material and Methods

### Seed Collections

Our study built on a unique collection of wheat material including three wild species *T. dicoccoides* (2n=28)*, T. boeoticum* (2n=14), *T. urartu* (2n=14) collected in the Near East and domesticated bread wheat *T. aestivum* (2n=42) collected in the Near East and North Germany (Suppl Fig. 1 and Suppl Table 1). We here refer collectively to these five wheat species and cultivars as wheat “genotypes”. More precisely, wild wheat accessions were sampled in a South-East region of Turkey, a region located in the Fertile Crescent and known to be the natural environment of these three wild wheat progenitors. Moreover, the region is considered to be a site of early domestication and cultivation of bread wheat *T. aestivum* (25). Our seed collections of the wild wheat represent two geographical populations of the wheat in central-eastern Turkish-Iraqi race (24). Seeds of the wild wheat were collected from one of the centers of massive stands in Karacadağ (provinces of Şanlıurfa and Diyarbakır) and Kartal-Karadağ (province of Gaziantep) in the South-East region of Turkey at different nearby fields in three years; 2004, 2005, and 2006 (Suppl Table1-Suppl Fig. 1) (24).

Seeds of domesticated wheat *T. aestivum* were obtained from a local farm in the same region where the wild wheat was collected. The *T. aestivum* genotype from Turkey is a winter wheat and local landrace of Kışlak, province of Hatay, and it has not been treated with chemicals by the farmer and only with a minimum amount of fertilizer (personal comm. by Nufel Gündüz, 2017). Also, we collected seeds from a modern winter wheat cultivar, Benchmark (IG Pflanzenzucht, Ismaning, Germany) originating from an experimental farm in Schleswig-Holstein, Germany. In contrast to the Turkish *T. aestivum*, this inbreed cultivar has been treated with chemicals during seed-production. Seeds of both *T. aestivum* genotypes were collected in 2017.

### Processing of the seed, leaves and roots

To ensure that we only isolate microbial DNA from the interior of seeds and tightly attached to the surface, we mildly surface semi-sterilized the seeds before DNA extraction. Seeds were surface-sterilized by shortly soaking them in TritonX 0.1%, 80% EtOH and 1.2% bleach followed by three washes with nuclease-free water. Three randomly selected samples from the wash-off water were also processed for sequencing as sterilization controls alongside the sterilized seeds.

Sterilized seeds were frozen by utilizing the Cryolys cooling unit and homogenized with a Precellys Evolution Tissue Homogenizer (Bertin Instruments, Montigny-le-Bretonneux, France). DNA was extracted from single seeds following a phenol-chloroform extraction protocol (see Suppl. Text). This method was developed from a previously established protocol for *Arabidopsis thaliana* (26), and here optimized to increase the efficiency of extraction of bacterial and fungal DNA from single seeds. Three randomly selected negative controls (i.e. blanks) of DNA extraction were also sequenced alongside the seed samples. Processing of DNA extracts and sequencing is described below.

We further addressed the colonization dynamics of the seedborne microorganisms in an *in-vitro* experiment where we germinated seeds under sterilized conditions in closed sterile jars to assess microbial diversity in leaves and roots. In brief, seeds from three wheat genotypes (*T. aestivum* from Turkey and Germany and *T. dicoccoides* from Turkey) (same seed populations used to characterize seedborne microbial diversity) were surface sterilized and germinated under sterile conditions with 16h light/8h dark cycles at 15°C (n=4-8 per population) in a climate chamber (Percival plant growth chambers, CLF PlantClimatics GmbH, Wertingen, Germany). In the sterile jars, plants were grown in a nutrient-rich PNM medium (see Suppl. Text). We let seedlings develop for two weeks, which corresponds to the emergence of the second leaf. About six centimeter of two leaves and multiple leaves and roots of two weeks old seedlings were harvested with sterile forceps and processed for DNA extraction. DNA extraction was performed using the PowerPlant Pro DNA Isolation Kit (Mo Bio Laboratories, Heidelberg, Germany) according to the manufacturer’s instructions.

### Transplant Soil Experiments

To address if domestication has entailed a change in the ability of plants to associate with microbial communities, we reciprocally transplanted domesticated and wild wheat (*T. aestivum* from Germany and Turkey and *T. dicoccoides* from Turkey) in a German agricultural soil and a natural soil from a region of the Fertile Crescent in Turkey. Both soil types were mixed with 5% peat and sifted with a sieve. We propagated seedlings from surface sterilized seeds in the two soil types in the climate chamber. We harvested leaves and roots as described above for the axenically propagated seedlings (n=6-8 per wheat accession-soil type combination). Additionally, three pots per soil type were filled with soil without plants and used as controls that were processed alongside samples. The position of each pot was being changed during the experiment to randomize any spatial bias. After two weeks, leaves and roots were harvested with sterile forceps and scissors and mildly washed with water, 1% PBS and 1% PBS + 0.02% Tween20 to remove loosely attached microbes and soil particles from the roots. Finally, samples were processed for DNA extraction. DNA extraction was performed using the PowerPlant Pro DNA Isolation Kit (MoBio Laboratories, Heidelberg, Germany) according to the manufacturer’s instructions.

### Sequencing of amplicons

The V5-V7 sequence of the bacterial 16S ribosomal RNA (16S rRNA gene) and a sequence of the fungal ribosomal internal transcribed spacer (ITS1) region were amplified using the primer combinations 799F-1192R and ITS1F-ITS2 to assess bacterial and fungal diversity, respectively (27), (28). Bacterial and fungal sequences were amplified with a two-step PCR protocol. In the first PCR step, interfering primers were utilized to enrich amplification of the 16S rRNA and preventing unintended co-amplification of the DNA. These interfering primers were originally developed by Agler and co-workers for microbial community analyses of *A. thaliana* (26). Here, we modified the interfering primers to target the corresponding wheat loci and changed the PCR protocol to optimize primer interference (Suppl. Text). In the second step of PCR, reverse primers barcoded with 12 base pair indexes and unique to each sample were used as barcodes to multiplex different samples in one sequencing run (Metabion International AG, Planegg, Germany). The primer setup used here was taken from Agler et al. 2016 (26). Three PCR replicates for each sample were used as technical replicates in each step and subsequently merged at the end of each PCR.

Finally, amplicon libraries were quantified fluorescently with the Invitrogen Qubit 3.0 Fluorometer (Thermo Fisher Scientific, Darmstadt, Germany). 16S and ITS amplicons were combined in equimolar concentrations in combined libraries. During DNA extraction as well as during library preparation, samples were randomized to prevent any possible batch effect. The combined libraries were paired-end sequenced for 600 cycles on an Illumina MiSeq machine at the sequencing facility of Max Planck Institute for Evolutionary Biology, Plön, Germany.

### Data Analysis

Raw reads were demultiplexed and converted into fastq files for downstream analysis using the bcl2fastq Conversion Software of Illumina (Illumina, bcl2fastq Conversion Software v2.20.0.422). We followed the QIIME2 version 2019.1 pipeline to preprocess and filter the fastq files (29). In brief, the conserved flanking regions of the ITS reads were trimmed with the q2-itsxpress plugin integrated into QIIME2 (30). Afterwards, the DADA2 software package also integrated into QIIME2 was used to correct and truncate sequences and filter chimeric reads for 16S reads (31). However, ITS reads were not truncated but only corrected and filtered as recommended by the q2-itsxpress plugin tutorial (https://forum.qiime2.org/t/q2-itsxpress-a-tutorial-on-a-qiime-2-plugin-to-trim-its-sequences/5780). After filtering and denoising, no fungal or bacterial features remained in the negative controls of DNA extraction and sterilization. Alpha diversity rarefaction plots for each sample confirm that a sufficient depth of coverage of the 16S and ITS datasets were achieved for both seeds and seedlings (Suppl Fig. 2). Further details regarding the amplification and sequencing is included in the supplementary materials and methods (see the Suppl Text).

For the taxonomic classification of 16S and ITS datasets, we used the Greengenes13.8 and UNITE 7.2 databases, respectively (32), (33). We utilized the q2-feature-classifier plugin of QIIME2 to extract the reference sequences from the databases and train the Naïve Bayes classifier (34). We extracted the target sequence of the 799F-1192R primer pairs from the Greengenes13.8 database. However, we did not extract the target sequences of ITS primers but used the full reference sequences as suggested by the q2-feature-classifier tutorial (https://docs.qiime2.org/2018.6/tutorials/feature-classifier/). Next, we trained the Naïve Bayes classifier based on the reference sequences and taxonomy. Finally, the resulting feature table was used to determine taxonomic relative abundances and for the subsequent statistical analyses of beta-diversity.

Downstream analyses were conducted with the “phyloseq”, “vegan”, “ampvis2” and “ggplot2” R packages or custom R scripts (35)), ((36)), ((37)), ((38)), ((39). Samples with fewer than 1000 reads for 16S and 200 reads for ITS were excluded from the resulting table. Moreover, taxonomically unassigned reads at the kingdom level and reads assigned to mitochondrial or other plant sequences were excluded for further analyses. Further information about the summary of the data before and after filtering is available in the supplementary material (Suppl Table 2). Before estimating alpha diversity indices, the samples were rarefied to even depth. Alpha diversity indices of the samples were estimated with Shannon diversity metrics and using observed number of features (i.e. richness) as a metrics. The significance of differences in diversity among wheat groups and pairwise multiple comparisons between wheat species were tested using a Kruskal-Wallis test (krus.test in R) and Conover’s test in the “PMCMR” package where we corrected the p-values with the “holm” correction method (40).

To compare the composition of communities and abundances of microbial taxa among different populations of wheat hosts, the counts from the feature tables were normalized by the cumNorm function in the “metagenomeSeq” package (41). We computed the Jaccard, Bray-Curtis and unweighted UniFrac distances to compare the structure of bacterial communities among/between samples. First metrics account for the absence/presence, second for both absence/presence and abundances whereas UniFrac metrics incorporates phylogenetic relatedness of bacterial communities into the calculation of distances. For fungal communities, we only used Bray-Curtis and Jaccard metrics. We did not report phylogeny-based metrics for the fungal data because sequence length variation in ITS may lead to erroneously inference of phylogeny. Beta diversity distance matrices were used for principal coordinate analysis (PCoA). Permutational multivariate analysis of variance analysis (PERMANOVA) was performed to test the significance of the effect of soil type and host type and their interactions in the microbial community composition (“adonis” function in the “vegan” package in R).

## Results

### Domestication of wheat has negligible effect on seedborne microbial diversity

To compare the diversity of seedborne microbiota between domesticated and wild wheat we firstly used the three wild wheat species *T. dicoccoides, T. urartu* and *T. boeoticum* and two genotypes of domesticated wheat *T. aestivum,* one landrace from Turkey and an inbred cultivar from Germany (Suppl Table 1). For *T. dicoccoides* we used genotypes from four different populations in South-East Turkey (Suppl Table 1). Our method of seed surface sterilization allowed us to assess microbial diversity exclusively of the seedborne microbiota of individual seeds. We extracted DNA from individual seeds and amplified microbial DNA using both bacterial and fungal specific primers (16S rRNA gene, V5-7 regions and ITS, ITS1 region, respectively) (Suppl Table 3 and Suppl Text).

Measures of alpha diversity (within sample diversity) show an overall low diversity of microbial taxa in the wheat seeds and notably a low diversity of seedborne fungal taxa compared to other plant tissues like leaves and roots. On average, in 58 samples, we found 68.7 bacterial features and 5.3 fungal features (corresponding to a Shannon Index of 2.6 and 0.8 for bacteria and fungi, respectively) (Fig. 1A and 1B). Notably, pairwise comparisons of alpha diversity among the different wheat genotypes showed no difference. Specifically, we did not observe any difference in microbial diversity associated with seeds of wild and domesticated wheat (Kruskal-Wallis test, Richness: p= 0.8029 and 0.1924; Shannon diversity: p= 0.6728 and 0.2530 for bacterial and fungal communities, respectively) or among the different wheat genotypes (Kruskal-Wallis test, Richness: p= 0.2421 and 0.4481; Shannon diversity: p= 0.6826 and 0.7191 for bacterial and fungal communities, respectively).

**Figure 1:**
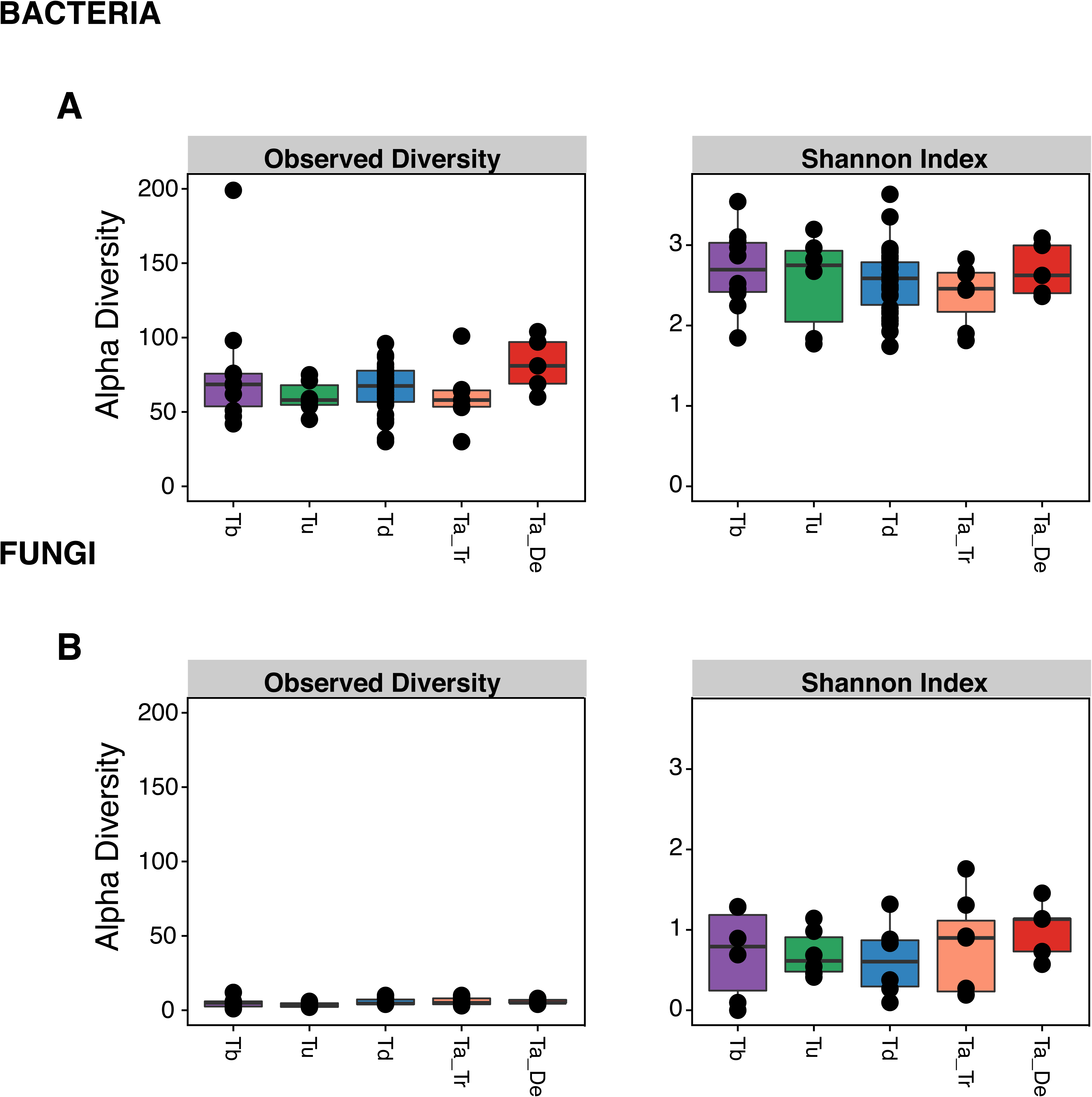
Alpha diversity in the seedborne microbiota of different wheat genotypes: Seeds of different wild and domesticated wheat harbour comparable microbial diversity. Alpha diversity of **A)** bacterial and **B)** fungal features in the seeds of different wheat. *T. boeoticum* (Tb), *T. urartu* (Tu), *T. dicoccoides* (Td), *T. aestivum* from Turkey (Ta_Tr) and *T. aestivum* from Germany (Ta_De). Each dot in the boxplots show the microbial diversity of a single seed. Pairwise comparisons of alpha diversity showed no significant difference between wheat genotypes (Conover’s Test: Richness p-values: 0.41-1.00 and 1.00, Shannon diversity p-values: 1.00 for bacteria and fungi, respectively).

Taken together, our estimates of alpha diversity in different wheat accessions and species suggest that domestication has not entailed a loss of diversity in the seedborne microbiota.

### Different composition of microbial communities associated with different wheat species

We next investigated the composition of microbial communities associated with the wheat seeds. Comparisons of beta-diversity (between sample variation), showed that the seedborne bacterial and fungal communities cluster independently of the wheat species (Suppl Fig. 3). Notably, the seedborne microbial communities of *T. aestivum* accessions from Germany and Turkey cluster together although the first represents a highly inbred modern cultivar and the second a local Turkish landrace.

We further characterized and compared the identity and abundances of microbial taxa. To this end, we aggregated the assigned taxonomy of each microbial feature to the family level. First, we assessed the distribution of major bacterial groups associated with seeds of the different wheat genotypes. We found a considerable variability among replicates of the same wheat genotype (on average 12.1% of bacterial features at the family level exist in all replicates of the same wheat genotype). Hereby, the wheat seed microbiota was mostly dominated by Proteobacteria and to lesser extent by Firmicutes, Actinobacteria and Bacteroidetes (Fig. 2A). These results are in accordance with previous studies of seed-associated bacteria of crop (e.g. maize, barley, rice) (42)(16), (42), (43)(16) and non crop plants (e.g. radish) (2), (44). However, at lower taxonomic levels, we observed differences in abundances of several microbes among the different wheat genotypes (Fig. 2A). For example, the Halomonadaceae family, including bacteria known to promote plant salt tolerance and growth (45), represent a substantial proportion of the bacterial community in the seeds of wild wheat (17.6-22.9%) but a small proportion of the domesticated wheat seed microbiome (5.2-7%).

**Figure 2:**
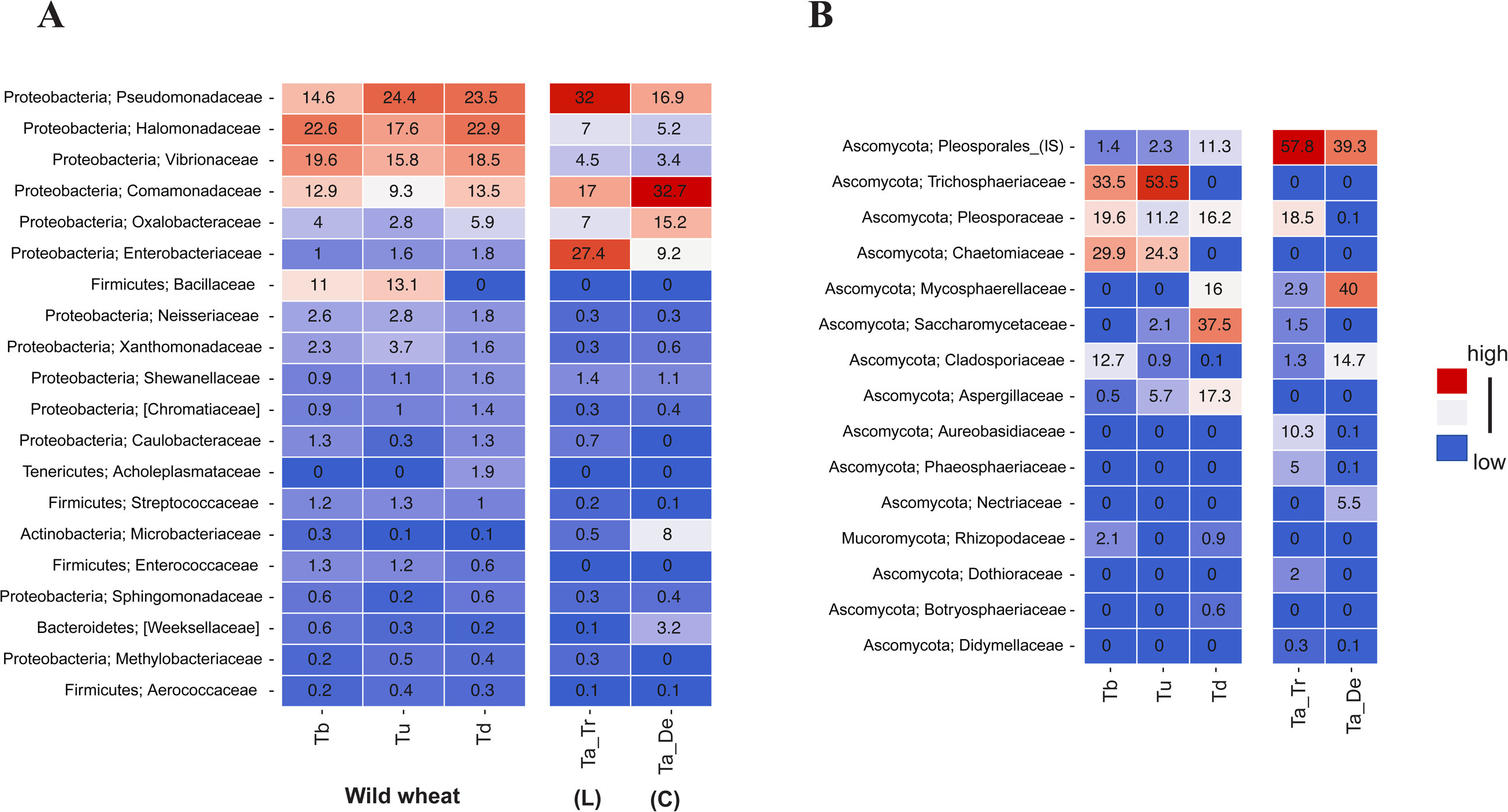
Composition of the seedborne microbiota across different wheat genotypes. Mean relative abundances of the most abundant **A)** twenty bacterial features, and **B)** fifteen most abundant fungal features at the family level in seeds of different wheat species. Color for each feature ranges from blue (minimum 0) to red with higher relative abundance values. (Abbreviations on figure, L: landrace, C: cultivar, IS: Incertae sedis taxa).

Among the fungal taxa, we also found a considerable variability among replicates of the same wheat genotype (on average 17.4% of fungal features at the family level exist in all replicates of the same wheat genotype). Fungal communities were dominated by Ascomycetes (Fig. 2B). Notably fungi in the order Pleosporales are abundant in the wheat seeds, including species of *Alternaria* that are highly abundant in seeds of *T. aestivum* and previously also shown to dominate wheat endophyte communities (46). Interestingly, Trichosphaeriaceae and Chaetomiceae were detected to be the most prevalent two fungal families in *T. boeoticum* (33.5% & 29.9%) and *T. urartu* (53.5% & 24.3%), were not detected in other wheat species.

In summary, while microbial alpha diversity is comparable among domesticated and wild wheat species, we report differences in the taxonomic composition of bacterial and fungal seedborne communities. These findings indicate that although domesticaion has a minor effect on the overall microbial community richness, it may have impacted the structure of these communities.

### Axenic seedlings of wild wheat are colonized by more diverse bacterial communities than domesticated wheat

We hypothesized that a proportion of the seedborne microbiota colonizes the plant seedling after seed germination. In order to compare microbial diversity and community composition of these early colonizers in domesticated and wild wheat, we set up an experiment using seeds of the German *T. aestivum* cultivar, Turkish *T. aestivum* landrace and the Turkish *T. dicoccoides* genotypes. We germinated surface-sterilized seeds and propagated these under sterile conditions. We harvested leaves and roots of the seedlings two weeks after seed germination, including a total of 32 plant samples consisting of roots and leaves (4-8 replicates per wheat and per tissue) and used these samples to profile bacterial and fungal communities.

Analyses of the bacterial microbiota revealed a total of 589 and 632 different bacterial features in leaves and roots (after filtering and rarefaction) (Suppl Table2). The analysis revealed that bacterial communities associated to the roots of *T. dicoccoides* are significantly more diverse compared to the communities associated with the Turkish and German *T. aestivum* genotypes (pairwise alpha diversity comparisons, Kruskal Wallis, Richness: p= 0.0023 and p= 0.0023; Shannon index: p= 0.0021; p= 0.0027, respectively). Additionally, the leaves of *T. dicoccoides* hosted more diverse bacterial communities compared to domesticated wheat from Turkey (Kruskal Wallis, Richness: p= 0.0280; Shannon index: p= 0.0066) (Fig. 3A). Taken together, significantly more diverse seedborne bacterial community was transmitting into leaves and roots in wild wheat compared to the domesticated wheat.

**Figure 3:**
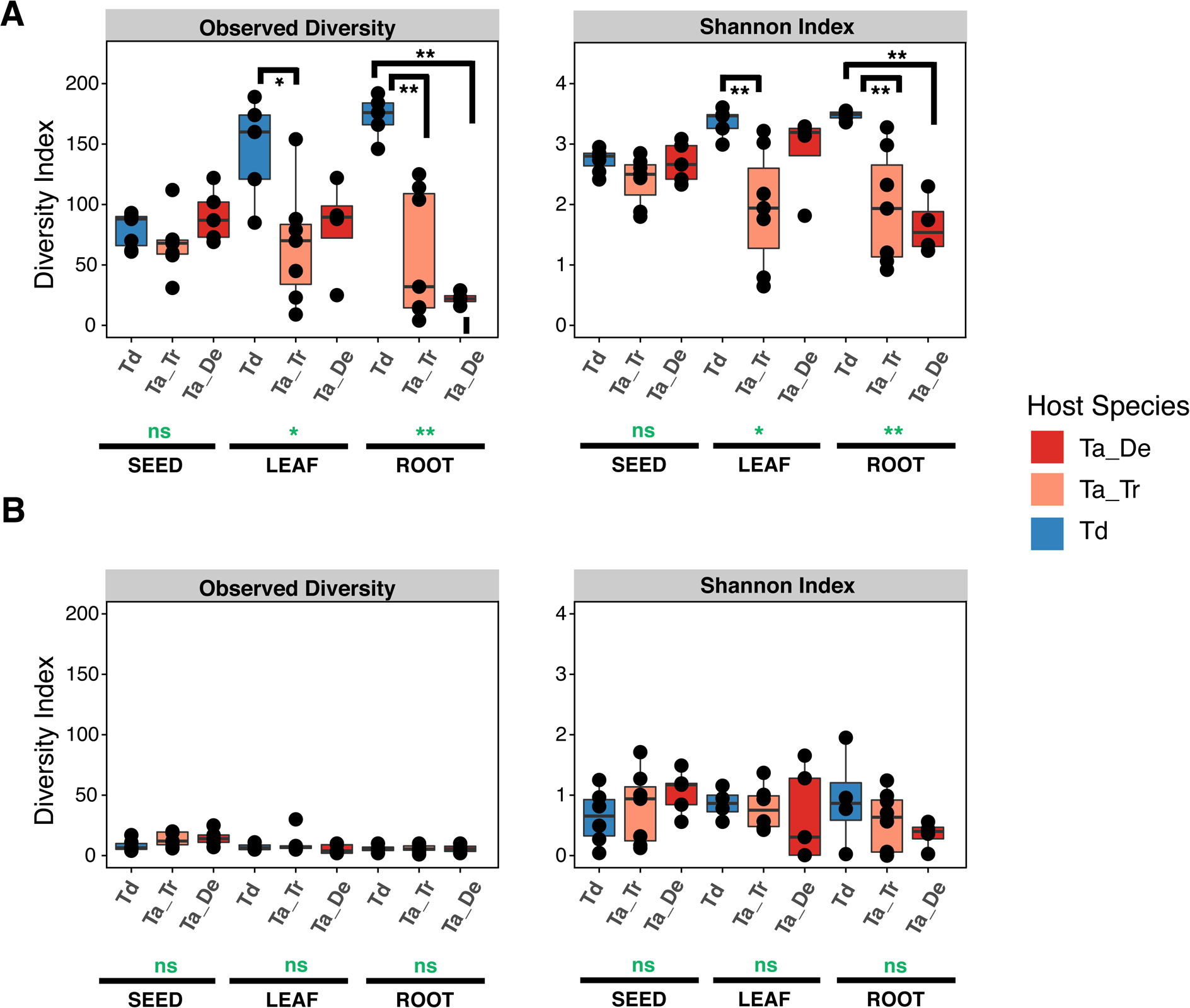
Significantly more diverse bacterial but not fungal communities are colonizing the wild wheat. Microbial diversity in different tissues of axenically-grown wheat: Alpha diversity of **A)** bacterial and **B)** fungal taxa in seeds, leaves and roots of the wild wheat *T. dicoccoides*, *T. aestivum* from Turkey and *T. aestivum* from Germany respectively. Global p-values for each tissue are shown in green and p-values of pairwise comparisons are in black color. (* < 0.05; ** <0.005; ns= non-significant)

Roots and leaves of the wheat seedlings were colonized by few fungal taxa, and we obtained in total only 98 and 74 unique fungal features in leaves and roots (after filtering and rarefaction). In contrast to the striking difference we observed for bacterial communities in wild and domesticated wheat, we observed no significant difference in the diversity of fungal colonizers suggesting that different processes determine the colonization of bacterial and fungal endophytes (Fig. 3B).

We next compared the identity and abundances of the microbial communities of seeds and seedlings. Overall, the same bacterial and fungal phyla were dominant in seeds as well as in leaves and roots, however we observed some significant shifts in microbial abundance (Fig. 4A). For example, Comamonadaceae, Halomonadaceaea, Vibronaceae and several other bacterial families enriched in seeds of both wild and domesticated wheat, did not colonize roots of the German *T. aestivum* accession. Furthermore, Paenibacillaceae was only present at very low abundance (0-0.1%) in seeds but a dominant colonizer of roots of *T. aestivum* from Germany (26.3%).

**Figure 4:**
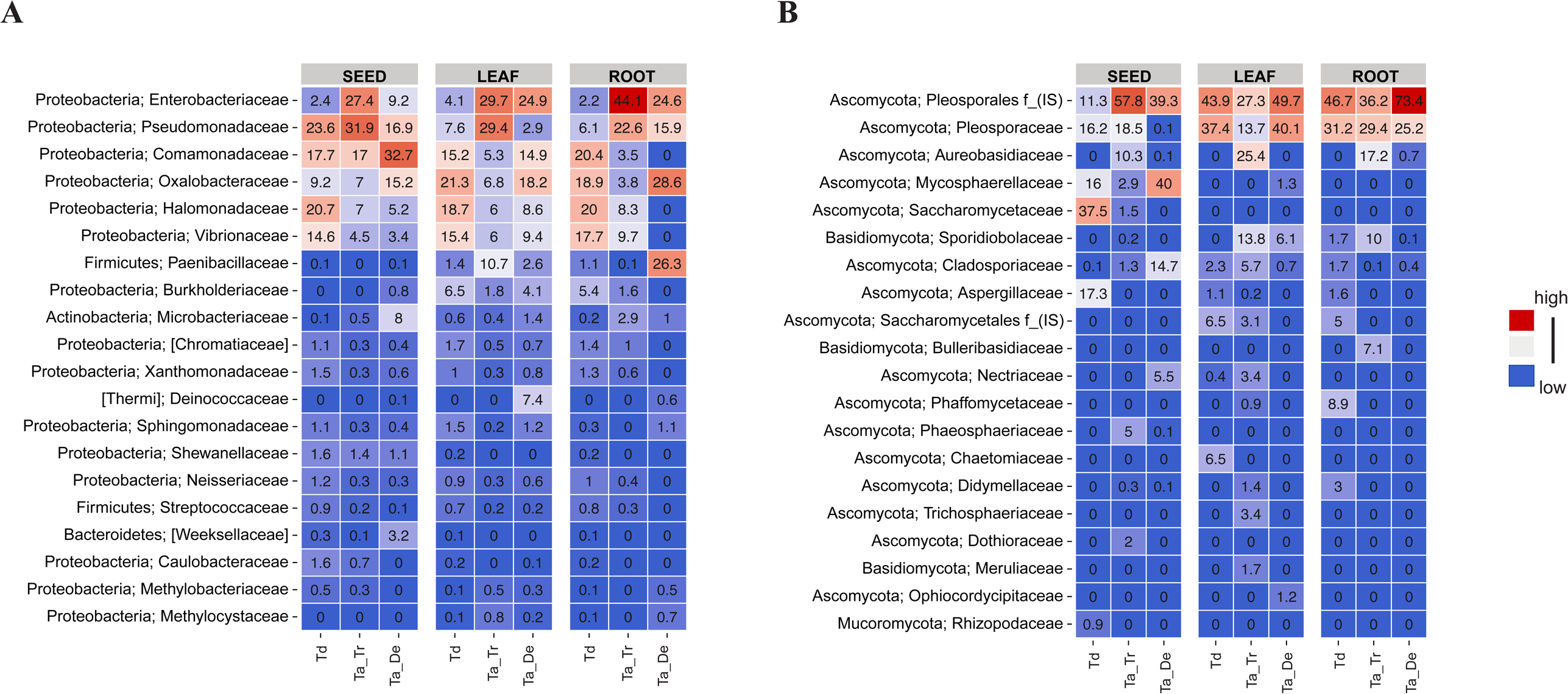
Composition of the microbiota in the axenic seedlings across different wheat genotypes. Mean relative abundances of the most abundant twenty **A)** bacterial and **B)** fungal features at the family level in seeds, leaves and roots of the German *T. aestivum* genotype Ta_De, the Turkish *T. aestivum* genotype Ta_Tr and the wild wheat *T. dicoccoides* genotype Td, respectively. Colors for each taxon illustrate relative abundance and ranges from blue (minimum 0) to red with higher relative abundance values. (Abbreviation, IS: Incertae sedis taxa).

Fungal ascomycete taxa were the most abundant in the wheat seedlings (Fig 4B). Notably, Pleosporales were abundant colonizers of seedlings of both wild and domesticated wheat from Turkey and Germany. Aureobasidiaceae were abundant only in the seedlings of *T. aestivum* from Turkey (25.4% in leaves and 17.2% in roots), but not in other wheat. Other abundant seedborne fungi were not detected in the leaves and roots of the wheat seedlings. For example, Mycospaerellaceae (40% in *T. aestivum* from Germany), Saccharomycetaceae (37.5% in *T. dicoccoides*); Aspergillaceae (17.3% in *T. dicoccoides*) found to be abundant in the seedborne communities were either absent or only present at low relative abundance in the seedlings. Together, these results demonstrate a difference in the assembly of seedborne bacterial, but not fungal communities in wild and domesticated wheat seedlings. Moreover, our results indicate that more diverse microbial communities are sustained in root and leaves of wild wheat seedlings compared to domesticated wheat.

### Axenically grown domesticated wheat seedlings assemble less homogeneous microbial communities

Our analyses of bacterial diversity in seeds and seedlings of *T. aestivum* and *T. dicoccoides* revealed a considerably fewer diversity in the replicates of *T. aestivum* seedlings (Fig. 3A). We further examined between-sample variation by computing Bray-Curtis and Jaccard and Unweighted UniFrac distance metrics (Fig. 5, and Suppl Fig. 3). Our results show that replicates of seedborne bacterial communities of the wild wheat and domesticated wheat from Turkey and Germany cluster together in the PCoAs (Fig. 5A and Suppl Fig. 3D-E). However, the bacterial colonizers of *T. dicoccoides* are distinct from the seedborne community and there is less variation among replicates of root and leaf communities. In contrast, in the domesticated *T. aestivum* wheat from Germany and Turkey, we observed more heterogeneous microbial communities associated with the leaves and roots (Fig. 5A). Pairwise comparisons of Bray-Curtis distances confirmed the distinct community composition of the wild and domesticated wheat: replicates of *T. dicoccoides* seedlings were colonized by notably more similar bacterial communities in comparison to replicates of *T. aestivum* seedlings (Fig. 5B).

**Figure 5:**
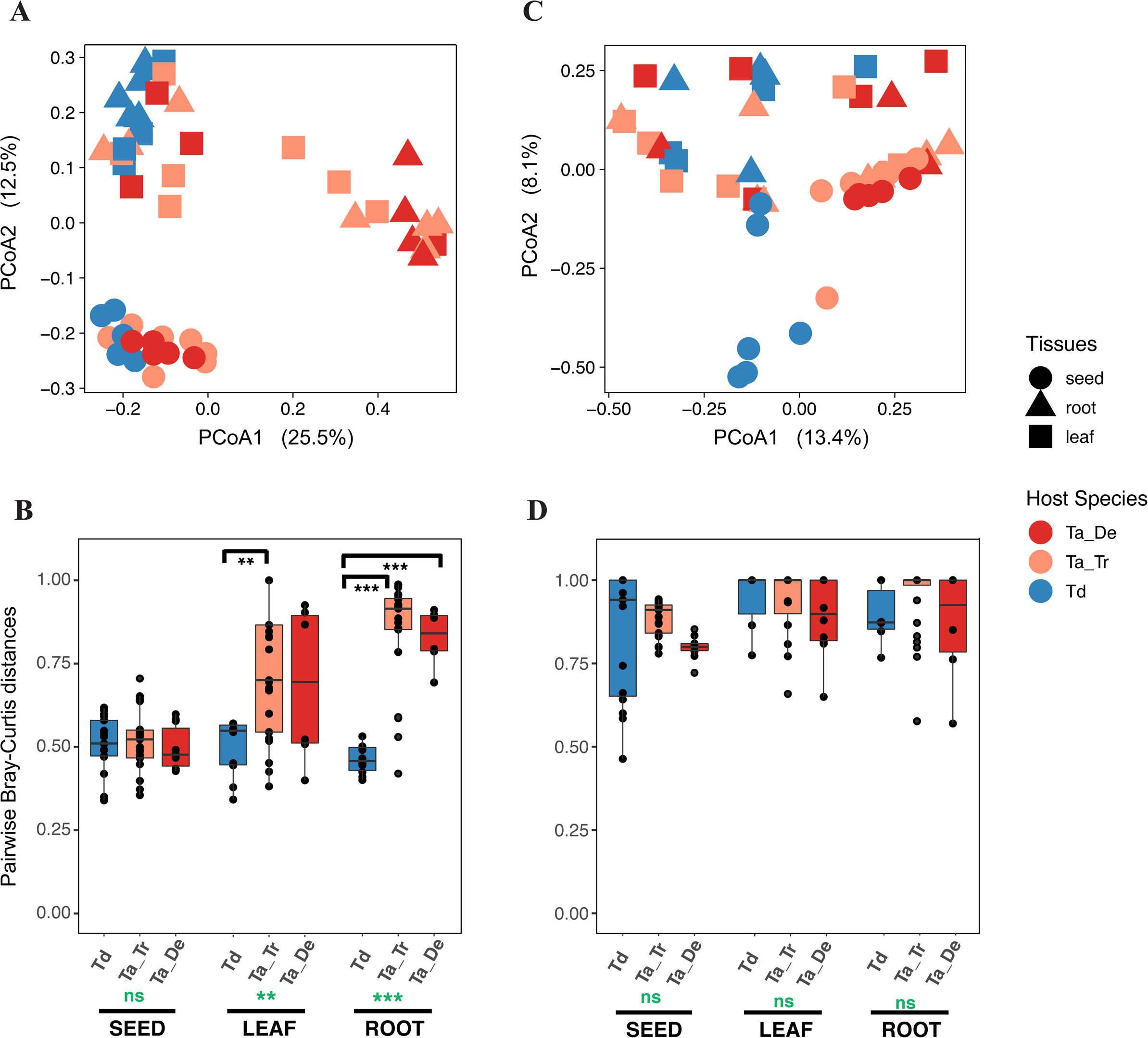
Wild wheat is colonized by more homogenous bacterial communities among different individual plants compared to domesticated wheat in roots. **A-B)** Bray-Curtis distance metrics based PCoAs of **A)** bacterial **C)** fungal communities of each seed, leaf and root sample from the axenic experiment. **B and D)** Estimated pairwise Bray-Curtis distances of **B)** bacterial and **D)** fungal communities in the replicates of each wheat genotype for each tissue. Global p-values for each tissue are shown in green and p-values of pairwise comparisons are in black color. (** <0.005; *** < 0.0005; ns= non-significant)

Additionally, we compared variability in fungal communities among seed and seedling replicates. In contrast to the seedborne bacterial communities of wheat, we found considerably more difference among replicates of seedborne fungi as well as colonizers of roots and leaves (Fig. 5C-D and Suppl Fig. 3D). Overall, these findings also support that different processes govern the assembly of bacterial and fungal communities in seeds and seedlings of wheat. We moreover conclude that wheat domestication and plant polyploidization did not entail a modification of the seedborne microbial diversity, but rather affected the diversity and composition of leaf and root colonizers.

### Domestication has not changed the assembly of soil-derived root microbiota

In their natural environment, plant seedlings are also colonized by microorganisms from the soil (47). To investigate if seedlings of wild and domesticated wheat assemble different microbial communities from soil, we set up an experiment with the same wheat genotypes used above. We germinated seeds and propagated seedlings of *T. dicoccoides* and *T. aestivum* in two different soils: an agricultural soil from Germany and a natural soil obtained from a location in the South-East region of Turkey close to the sampling site where the wild wheat accessions were obtained. We propagated the three wheat genotypes (*T. dicoccoides* from Turkey and *T. aestivum* from Turkey and from Germany) independently in the agricultural and the natural soil (6-8 replicate plants per wheat-soil combination). The seeds used here were surface sterilized as in the experiments described above.

Our results reveal that soil type rather than plant genotype is a main determinant of bacterial community structure in roots of the wheat seedlings (using a PERMANOVA test, explained by 61.38% of the between-sample variation; p= 0.001) (Fig. 6A-B). However, for the leaf-associated microbial communities, the wheat genotypes explain a significant proportion of the bacterial diversity (for soil type 13.54%; p= 0.001, for wheat accession 6.09%; p= 0.020 and for the interaction of soil and wheat accession 5.31%; p= 0.066) (Fig. 6 and Suppl Fig. 12).

**Figure 6:**
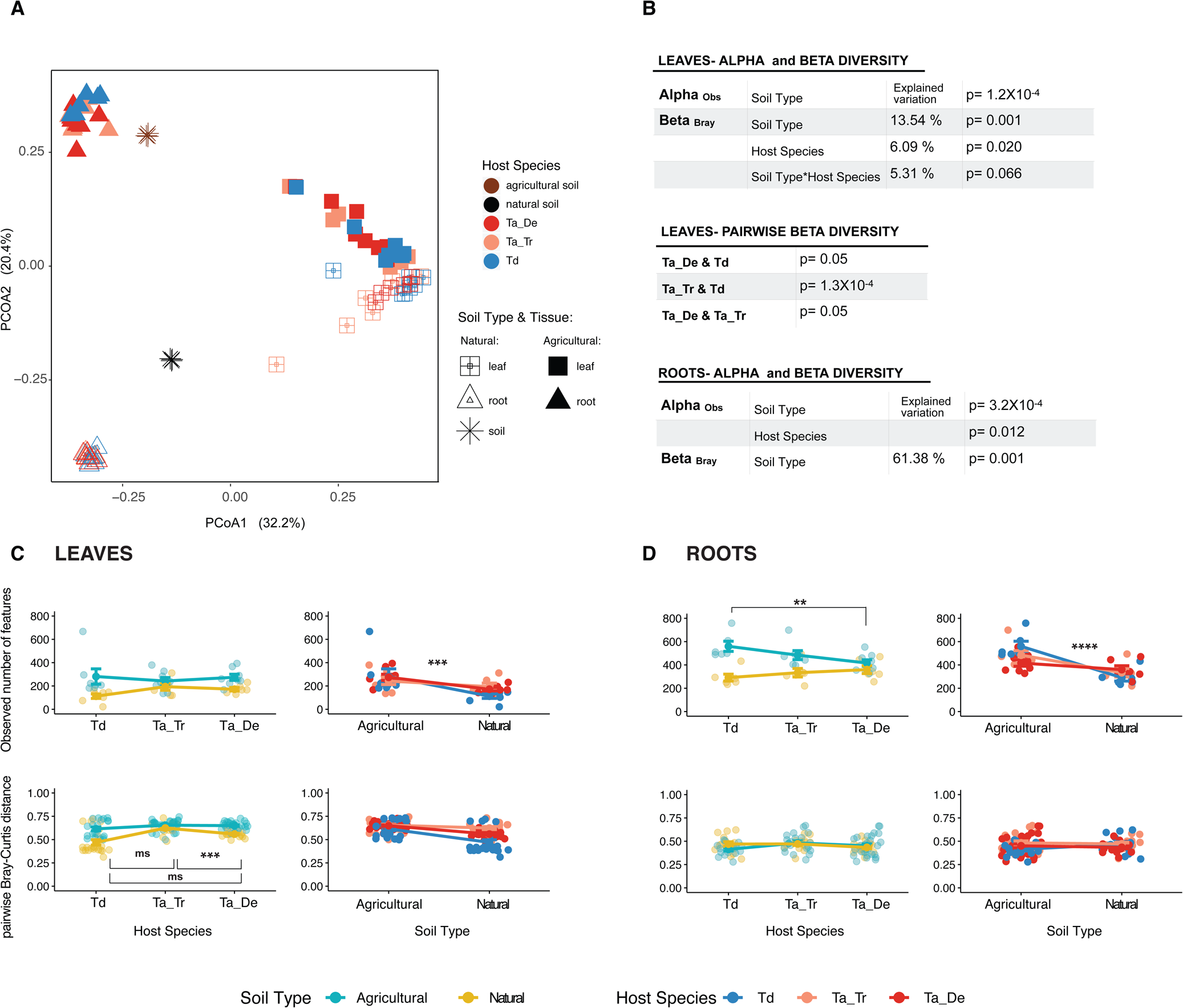
Soil type in roots, both soil type and wheat genotype in leaves are the determinants of the bacterial community. **A)** Bray-Curtis distances based PCoA of bacterial communities of plants grown in soil **B)** Summary statistics for the beta-diversity and significant alpha diversity comparisons of the bacterial communities **C-D)** Interaction plots showing alpha and beta diversity comparisons of bacterial communities in **C)** leaves and **D)** roots of different wheat grown in diffent soil types.

In general, plants grown in the agricultural soil were colonized by more diverse bacterial communities in leaves and roots (Fig. 6C and 6D). In contrast to the axenic experiment, we did not detect a difference in alpha diversity between leaves of wild and domesticated wheat when seedlings were propagated in the soil. However, in accordance with our observations from the axenic experiment, we observe less variation between replicates of *T. dicoccoides* compared to domesticated *T. aestivum* when the seedlings were propagated in the natural soil. On the other hand, we observe a significant difference in alpha diversity between microbial communities of *T. dicoccoides* and the German accession of *T. aestivum* propagated in the agricultural soil. Hereby, the bacterial diversity is higher in roots of the wild wheat when compared to the domesticated wheat. We speculate that the higher diversity of the bacterial communities associated with *T. dicoccoides* in the foreign soil originates from novel plant-microbe interactions.

The composition of fungal communities was assessed only in roots of the three wheat genotypes (Suppl Fig. 10 and Suppl Fig. 11). Our results show that soil type but not wheat genotype is also the main determinant of fungal community structure associated with the wheat roots (PERMANOVA test, %8.52 of the between-sample variation; p= 0.001). Notably, when the seedlings were propagated in the natural soil, they were colonized by significantly more diverse fungal communities (p= 3.71 × 10^−6^ and p= 8.90 × 10^−4^ for richness and Shannon Index, respectively) (Suppl Fig. 11A-B). Moreover, based on analyses of pairwise Bray Curtis distances we found that fungal communities are more similar among replicates when seedlings were propagated in the natural soil compared to the agricultural soil (Suppl Fig. 11C). Also, the wild wheat were colonized by less homogenous fungal communities compared to the domesticated wheat in both soil types. Taken together, the differences in diversity and community composition of fungal wheat colonizers had little effect of the wheat genotype, but a significant effect of the soil type.

We next compared the identity and abundances of microbial taxa in the seedlings of the wheat seedlings propagated in soil. Clearly, roots and leaves exhibited distinct bacterial composition when compared to the bacterial communities of the bulk soil implying specificity related to plant colonization (Fig. 6A and Suppl Fig. 13A). However, in general microbial composition was similar among the three different wheat genotypes when grown in the same soil type (Suppl Fig. 13A). Basically, roots were dominated by two bacterial families; Oxalobacteraceae (18.6-26.5%), Streptomycetaceae (27-32.6%) (Suppl Fig. 13A) where Streptomycetaceae were the dominant colonizer of roots when seedlings were growing in the agricultural soil (40.2-49.7%) (Suppl Fig. 13C). On the other hand, leaves were colonized by other bacterial families: Oxalobacteraceae (12.2-22.2%), Comamonadaceae (7.4-19.9%), Rhizobiaceae (12.1-21.3%), Halomonadaceae (13.6-15.9%), Vibronaceae (8.4-14.7%). Notably, the two bacterial families Halomonadaceae and Vibronaceae were highly abundant phyllosphere colonizers in leaves propagated in both soil types (Suppl Fig. 13B), although they were not detected in the bulk soil or roots (0-0.1%). On the other hand, these bacteria were also prevalent members of the seedborne leaf community suggesting that they may have originated from the seeds (Fig. 4A).

Also the fungal communities were similar among the different wheat genotypes when grown in the same soil type (Suppl Fig. 14). The most prevalent member of seedborne fungal communities, Pleosporaceae, were still detectable in the roots of plants propagated in the natural soil (10.8-16.6%). However, plants grown in the agricultural soil were mostly colonized by the fungal taxa in the family Pseudeurotiaceae (28.4-33.9%).

## Discussion

Plant domestication has entailed a significant loss of genetic diversity, as well as physiological and anatomical changes in the selected species (e.g. 18). In this study we have addressed to which extent domestication has changed the seedborne microbiota composition and the potential of wheat to assemble environmental (i.e. soil) microbial communities. We combined experimental assays with microbial profiling of a unique collection of wheat genotypes from a region in the Fertile Crescent, the center of origin of wheat. We specifically optimized our protocols of DNA extraction and microbial amplification to characterize the microbial community of individual seeds.

Comparing microbial communities associated with seeds, we showed significant differences in the composition of the bacterial communities in wild and domesticated wheat. This finding is consistent with previous studies, which have also revealed some differences in microbial community composition of domesticated plants and their wild relatives (48), (47), (49). Leff and co-workers investigated the impact of domestication on seedborne microbial communities of different sunflower accessions. Consistent with our results, they did not detect differences in fungal diversity between modern and wild sunflower and they demonstrated a minimal vertical transmission of fungal endophytes from seeds to roots of the developing seedling (30).

A growing body of evidence suggests that different layers of the plant immune system play significant roles in shaping the plant microbiota (50). Plants receptors that are involved in microbial recognition and “management” involve Pathogen Recognition Receptors (PRRs) and Nucleotide-Binding Leucine-rich repeat Receptor (NB-LRR) proteins (51). Wheat domestication and polyploidization may have conferred a change in the composition and diversity of these immune-related proteins and thereby indirectly impacted the assembly of the wheat microbiome. Comparative genome studies of domesticated and wild wheat have identified signatures of domestication in the *T. aestivum* genome, which may correlate with microbial community assembly, including changes in the repertoire of genes involved in signaling, hormone production and metal accumulation (52). We speculate that different microbial community composition in German and Turkish *T. aestivum*, and the wild relative *T. dicoccoides* reflect genetic differences among these wheat genotypes. We note that such changes notably have impacted the composition of bacterial communities and to a much lesser extent fungal communities. Interestingly, we observe little difference among seed and soil-derived fungal communities of the wild and domesticated wheat species. This suggests that different traits and mechanisms are responsible for the assembly of bacterial and fungal plant-associated communities.

The seedborne microbiome of wheat has previously been investigated primarily with culture-dependent methods (15), (46). Robinson and colleagues characterized seedborne microbial community in roots and shoots of the axenically grown seedlings by isolation and cultivation of microbiota (12). This study allowed them to characterize only eight bacterial taxa that could be defined at the genus level. Olfek-Lalzar also studied fungal endophytes in seeds of wild and domesticated wheat also using a culture-dependent method. They were able to isolate and identify 31 OTUs from 100 seeds; of these fungi more than half occurred only one time. In our study based on culture-independent methods and optimized molecular protocols, we were able to identify significantly higher diversity of bacteria and fungi than previously reported in wheat seeds (Suppl Table 2). Our data thereby provides an extended resource for further research of the wheat “core” microbiota, as well as species-specific microbial partners of modern and wild wheat.

An underlying assumption in our research of the seedborne microbiota is that a significant proportion of these microorganisms later represent endophytes in the wheat plant. To qualitatively and quantitatively characterize the early seedling colonizers we propagated seedlings under sterile conditions and assessed microbial diversity in leaves and roots of the young seedlings. A striking finding from this experiment was 1) a higher diversity of bacterial colonizers in the wild wheat *T. dicoccoides* compared to *T. aestivum*, and 2) an inconsistent community composition among replicates in *T. aestivum* seedlings. We speculate that stochastic processes (e.g. priority effects) play a stronger role in *T. aestivum* and result in the dominance of random bacterial taxa in different replicates. We see this effect both in the Turkish landrace of *T. aestivum* as well as in the inbred German wheat cultivar Benchmark, suggesting that the effect is consistent among different *T. aestivum* genotypes. We speculate that domestication may have entailed less selective constraints on plant traits that contribute to microbial assembly. Consequently, non-deterministic events could play a larger role in the assembly of microbial communities associated with domesticated wheat. Nonetheless, we note that the 16S and ITS data only allow a low-resolution analysis of microbial diversity and a more comprehensive investigation of plant colonization will require strain specific markers or specifically labeled strains.

Domestication and plant breeding have involved strong artificial selection of desired crop traits. For several domesticated species it is demonstrated that a negative consequence of domestication is a severe loss of genetic variation and an accumulation of deleterious mutations (53), (54), (21). These “domestication costs” may have reduced local adaptation of crop plants to their environment, including the local environmental microbiota. We addressed signatures of plant adaptation to the soil microbiota in the Turkish *T. dicoccoides* using local soil from Turkey. We compared plant microbial diversity of this local combination to a foreign combination of *T. dicoccoides* in a German soil. Moreover, we also assessed microbial diversity of two *T. aestivum* wheat in two soils. Although the two soils comprise different geochemical properties, they were comparable in their overall microbial diversity. Notably, we find that soil rather than plant genotype determines the composition of root associated bacterial communities suggesting that the “plant-selected” proportion of the soil-derived microbiota overall is small. Interestingly, we observe a stronger effect of the plant genotype on the bacterial phyllosphere community than on the root-associated bacterial communities. We speculate that the phyllosphere imply a strong selection on the associated microbiota. More detailed analyses of microbial diversity e.g. based on metagenome sequencing or microbial population genomic data is needed to study plant-microbe co-adaptation.

In conclusion, the present study provides new insights into the microbial community composition and colonization of domesticated and wild wheat. Our findings indicate an increased role of chance events and priority effects on seedling colonization. We moreover speculate that the difference between wild and domesticated wheat reflect changes in the plant immune system conferred by artificial selection and polyploidization during domestication and crop improvement. Future crop breeding strategies should account for microbial diversity and the ability of crops to assemble and maintain beneficial microbial communities. Such efforts will rely on research of plant-microbial co-adaptation and the underlying mechanisms that determine microbial plant colonization.

## Acknowledgements

The authors would like to thank Eric Kemen for helpful discussions, employees of the experimental farm Hohenschulen of Kiel University and Doreen Landermann for technical assistance. The authors also thank the organizers of the EMBO workshop „Plant Microbiota 2017“ for their input. This work was funded by the DFG Collaborative Research Centre (CRC) 1182 “Origin and Function of Metaorganisms” and the International Max Planck Research School for Evolutionary Biology. The authors declare no competing interests.

## Author Contributions

The study was conceived and designed by EÖ, EHS and TD. Seed collections and soil from Turkey were provided by HÖ and US. The experiments were performed by EÖ and MAH. Amplicon sequencing was performed by SK. Data analysis was conducted by EÖ and TD. The manuscript was written by EÖ and EHS. All authors read the manuscript.

## List of Supplementary Material

### Supplementary Text

**Suppl Table 1: Detailed information about the wheat collections**

**Suppl Table 2: Summary of the sampling and sequencing data**

“b.r.” denotes number of microbial features before filtering and rarefying while “a.r.” denotes number of features after rarefaction.

**Suppl Table 3: Sequence of the V5-7 and ITS1 primers used for amplification and sequencing**

**Suppl Figure 1: Location of the seed collections**

**A)** Samples were collected from Turkey and Germany **B)** Location of fields from South-East region of Turkey and North Germany where seeds were collected. Zoomed version of the locations was depicted to make visualization easier.

**Supp Figure 2: Alpha rarefaction curves for bacterial and fungal communities of each sample collection of each experiment A)** 16S seed samples **B)** ITS seed samples **C)** 16S axenic leaves and roots **D)** ITS axenic leaves and roots **E)** 16S leaf samples grown in soil and **F)** ITS root samples grown in soil **G)** ITS roots grown in the soil

**Suppl Figure 3: UniFrac and Jaccard distances based PCoAs of the microbial community in seeds and axenic seedlings A)** Bray-Curtis PCoA-seed-associated bacteria and fungi

**B)** Jaccard PCoA-seed-associated bacteria and fungi **C)** Unweighted UniFrac PCoA- seed-associated fungi **D)** Jaccard PCoA-axenic seedling-associated bacteria and fungi **E)** Unweighted UniFrac PCoA- axenic seedling-associated bacteria

**Suppl Figure 4: Pairwise Bray-Curtis distances of the bacterial communities in the replicates of seeds**

Each dot shows pairwise distance between two replicates. Red boxplots show distances of the microbial communities between samples of the same wheat genotype. Black boxplots show distances of the bacterial communities between samples of two corresponding wheat genotypes indicated.

**Suppl Figure 5: Pairwise Unweighted UniFrac distances of the bacterial communities oft he replicates of seeds**

Each dot shows pairwise distance between two replicates. Red boxplots show distances of the microbial communities between samples of the same wheat genotype. Black boxplots show distances of the bacterial communities between samples of two corresponding wheat genotypes indicated.

**Suppl Figure 6: Pairwise Bray-Curtis distances of the bacterial communities in the replicates of seeds and axenic seedlings**

Each dot shows pairwise distance between two replicates. Red boxplots show distances of the microbial communities between samples of the same wheat genotype. Black boxplots show distances of the bacterial communities between samples of two corresponding wheat genotypes indicated.

**Suppl Figure 7: Pairwise Unweighted UniFrac distances of the bacterial communities in the replicates of seeds and axenic seedlings**

Each dot shows pairwise distance between two replicates. Red boxplots show distances of the microbial communities between samples of the same wheat genotype. Black boxplots show distances of the bacterial communities between samples of two corresponding wheat genotypes indicated.

**Suppl Figure 8: Pairwise Bray-Curtis distances of the fungal communities in the replicates of seeds**

Each dot shows pairwise distance between two replicates. Red boxplots show distances of the microbial communities between samples of the same wheat genotype. Black boxplots show distances of the bacterial communities between samples of two corresponding wheat genotypes indicated.

**Suppl Figure 9: Pairwise Bray-Curtis distances of the fungal communities in the replicates of seeds and axenic seedlings**

Each dot shows pairwise distance between two replicates. Red boxplots show distances of the microbial communities between samples of the same wheat genotype. Black boxplots show distances of the bacterial communities between samples of two corresponding wheat genotypes indicated.

**Suppl Figure 10: Bray-Curtis distances based PCoA of the fungal communities of roots grown in the agricultural and natural soil**

Data based on ITS amplicon data from *Triticum aestivum* (Ta_De and Ta_Tr) and *Triticum dicoccoides* (Td).

**Suppl Figure 11: Interaction plots of fungal communities of root samples grown in the soil from agriculture and wild**

Data based on ITS amplicon data from *Triticum aestivum* (Ta_De and Ta_Tr) and *Triticum dicoccoides* (Td).

**Suppl Figure 12: Interaction plots showing Shannon Index estimates of bacterial communities in seedlings grown in soil**

Data based on 16S amplicon data from *Triticum aestivum* (Ta_De and Ta_Tr) and *Triticum dicoccoides* (Td).

**Suppl Figure 13: Composition of the bacterial community in seedlings colonized by the soil-derived microbiota A)** Bacterial community in different tissues **B)** Leaf-associated bacterial community in plants grown in agricultural and natural soil **C)** Root-associated bacterial community in plants grown in agricultural and natural soil

**Suppl Figure 14: Composition of the fungal community in roots colonized by the soil-derived microbiota**

Data based on mean read abundances of fungal features. Analyses based on ITS amplicon data from *Triticum aestivum* (Ta_De and Ta_Tr) and *Triticum dicoccoides* (Td). (IS) stands for the “Incertae sedis” taxa.

