## Supplementary Materials for "Higher stochasticity of microbiota composition in seedlings of domesticated wheat compared to wild wheat"

#### Supplementary Methods

##### Origin of the wheat material

Seed collections of *Triticum dicoccoides, Triticum boeoticum* and *Triticum urartu* were collected from different locations in South- East Turkey in previous years (2004, 2005, 2006) (Supp. Table 1). Seeds of *T. aestivum* from Turkey was collected by the farmer from a field in Kışlak, Antakya in 2017 (Supp. Table 1) and stored in a dry and cold room. We obtained these seeds in the same year from the farmer and we stored them at 4 °C until further usage. Seeds of *T. aestivum* from Germany (Benchmark cv.) were collected from the experimental farm of the Kiel University in Hohenschulen, Schleswig-Holstein in 2017. All the seeds in the collections were stored at 4 °C until further usage and processed all the seeds in the collections in 2018.

##### Surface sterilization of seeds

Seeds of *Triticum aestivum*, *Triticum dicoccoides, Triticum boeoticum* and *Triticum urartu* were washed with sterile miliQ water. Later, each seed was treated shortly with 0.1% TritonX and washed with water, then 80% EtOH and washed with water and lastly 1.2 % Sodium Hypochlorite (NaClO) and finally washed with water three times. Wash-off water from some seeds were kept and sequenced. Some seeds were immediately processed for DNA extraction while others were grown in sterile jars.

#### Axenic propagation of leaves and roots

After sterilization, seeds were directly transferred into the plant minimal medium (PNM). 1L of PNM medium is composed 1 mL of of 500 mM Potassium Nitrate, 1mL of KH2PO4 stock solution (5g/ 100mL), 1 mL of K2HPO4 stock solution (2.5 g/100mL), 1mL of 2M Magnesium Sulfate (MgSO4 X 7 H2O), 1 mL of 200 mM Calcium Nitrate, 2.5 mL of Fe-EDTA and 1mL of Sodium Chloride stock solution (2.5 g/100 mL). The pH of the solution was adjusted to 6.2 and the medium was solidified by adding Gelrite. Finally, the medium was autoclaved at 121 $^{o}$C for 15 minutes. After autoclaving, 10 mL 1M filtered MES was added to the medium. The medium was transferred into sterile jars and after it was solidified, sterilized seeds were transferred into the medium. Seeds were germinated in sterile jars in a climate chamber (Percival plant growth chambers, CLF PlantClimatics GmbH, Wertingen, Germany). Light period of 16/8 hours, temperature 15 $^{o}$C) for two weeks corresponding to the second leaf growth stage.

#### DNA Extraction from seeds and seedling material

We followed a similar method to the Phenol-Chloroform extraction method customized by Agler et al, 2016 (1). For DNA extractions, seed, leaf and root materials was processed as follows: Single seeds were transferred into 7mL tubes containing sterilized zirconium beads. Approximately six cm of leaves and six cm of multiple roots were used for DNA extraction. We crushed samples with Bertin Precellys Instrument (Bertin Instruments, Montigny-le-Bretonneux, France) pre-cooled with liquid nitrogen at 7500 rpm for 2 x 30 seconds with an in-between pause of 15 seconds. To the crushed samples, we added 0.6 mL of DNA extraction buffer (0.5% SDS, 50 mM TRIS buffer at pH 8, 200 mM NaCl, 2mM EDTA). Next, we added 50 µg/mL of Lyzozyme and 10 µg/mL of Proteinase K and incubated the samples for 45 min at 37$^{o}$C. After incubation, samples were transferred into new tubes containing 0.1 mm zirconium and 0.5mm glass beads and beat-treated using Bertin Precellys (Bertin Instruments, Montigny-le-Bretonneux, France) at 6500 rpm for 2 x 30 seconds with a 15 second pause. 10 µg/mL RNAse was added to each sample before incubation at 37°C for 45 minutes. We next centrifuged the samples at 13000 rpm for 2 minutes to remove beads and non-homogenized tissue. The nucleic acids were cleaned up with phenol/chloroform/isoamyl alcohol (25:24:1) for three times, and chloroform/isoamyl alcohol (24:1) one time, then precipitated by adding 1/10th of the sample volume of 3 M sodium acetate and 2.5x volume 100% ethanol. Samples were then centrifuged at 4°C at 13000 rpm for 30 minutes. Afterwards, the pellet was washed with 70% ethanol. Finally, pellets were washed in 100 µL of sterile water and stored at -20 °C until further usage. DNA extracted from seeds was treated with 1:1 with 20% Chelex-100 for 30 minutes to remove potential PCR inhibitors. Harvested leaves and roots were processed with the PowerSoil DNA kit (Mo Bio Laboratories, Heidelberg, Germany) according to the manufacturer's instructions (See Material and Methods).

**DNA library preparation**

We prepared libraries for replicates of indvidual seed as well as leaf and root samples, respectively. In addition, we included three negative controls of DNA extraction and three wash-off controls from the surface sterilization to the libraries.

PCR amplification was performed in two steps (Suppl. Table 1 and 2). For 16S, we used the interfering primer approach developed by Agler and co-workers for *Arabidopsis thaliana* (Suppl. Table 3). However, we modified the primers according to the wheat genome (2). In the first step, universal primers primers Bacteria V5/V6/V7: B799F/ B1192R and for fungi ITS1: ITS1F/ITS2 were used to amplify the targeted regions (Suppl Table 3). Interfering primers for the 16S region were added in this step. Triplicates of each reaction were performed for both first and second steps. First PCR was run for 15 and 20 cycles for 16S and ITS, respectively. After the first reactions, three replicates were combined. 10 µL of each PCR product was cleaned with 0.5 µL Antarctic phosphatase and 0.5 µL Exonuclease I in 1.22 µL Antarctic phosphatase buffer (New England Biolabs GMBH, Frankfurt, Germany) at 37 °C for 30 minutes followed by 80 °C for 15 min. The product of the cleaned samples was used as a template for the second PCR in which unique barcodes for sequencing of 12bp length were added as well. Second PCRs were run for 20 and 15 cycles for 16S and ITS, respectively including in total 35 cycles for both 16S and ITS reactions.

**Table 1: PCR conditions for the V5-7 amplification**

| | | | | |
|---|---|---|---|---|
| **V5-V7** | | | | |
| **1st PCR:** | | **1st PCR conditions:** | | |
| 10 µL | Phusion Master Mix | 98°C | 30s | |
| 1 µL | Forward Primer | **98°C** | **10s** | |
| 1 µL | Reverse Primer | **55°C** | **45s** | **X15 cycles** |
| 2 µL | Interfering Forward Primer | **72°C** | **30s** | |
| 2 µL | Interfering Reverse Primer | 72°C | 5m | |
| 0.5 µL | DMSO | | | |
| 3 µL | DNA | | | |
| 0.5 µL | H2O | | | |
| Clean-up with ExoI and Antartic Phosphatase | 10 µL oft he 1st PCR product is cleaned-up | 37°C<br>80°C | 30m<br>15m | |
| **2nd PCR:** | | **2nd PCR conditions:** | | |
| 10 µL | Phusion Master | 98°C | 30s | |

| | | | | |
|---|---|---|---|---|
| | Mix | | | |
| 1 µL | Forward Primer | **98°C** | 10s | |
| 1 µL | Barcoded Reverse Primer | **55°C** | 45s | **X20 cycles** |
| 0.5 µL | DMSO | **72°C** | 30s | |
| 3 µL | Clean-up product | 72°C | 5m | |
| 4.5 µL | H2O | | | |

**Table2: PCR conditions for the ITS1-ITS2 amplification**

| | | | | |
|---|---|---|---|---|
| **ITS1F-ITS2** | | | | |
| **1st PCR:** | | **1st PCR conditions:** | | |
| 10 µL | Phusion Master Mix | 98°C | 30s | |
| 1 µL | Forward Primer | **98°C** | **10s** | |
| 1 µL | Reverse Primer | **55°C** | **30s** | **X20 cycles** |
| 0.5 µL | DMSO | **72°C** | **30s** | |
| 1 µL | DNA | 72°C | 5m | |
| 6.5 µL | H2O | | | |
| Clean-up with ExoI and Antartic | 10 µL of the 1st PCR product is | 37°C<br>80°C | 30m<br>15m | |

| | | | | |
|---|---|---|---|---|
| Phosphatase | cleaned-up | | | |
| **2nd PCR:** | | **2nd PCR conditions:** | | |
| 10 µL | Phusion Master Mix | 98°C | 30s | |
| 1 µL | Forward Primer | **98°C** | **10s** | |
| 1 µL | Barcoded Reverse Primer | **55°C** | **30s** | **X15 cycles** |
| 0.5 µL | DMSO | **72°C** | **30s** | |
| 1 µL | Clean-up product | 72°C | 5m | |
| 6.5 µL | H2O | | | |

Finally, PCR products of ITS and 16S amplifications were sub-pooled in equal concentrations where the concentration of each sample was estimated using the software of gel visualizer (BIO RAD, Image Lab$^{TM}$ software 5.2.1). Subpools of 16S amplicons were run on the gel and bacterial DNA was cut out of the gel. DNA concentration of the subpools were quantified with Qubit. Then, we combined subpools in equal concentrations into final pools. Finally, we quantified the final concentration of the final pool with Qubit 3.0 Fluorometer (Thermo Fisher Scientific, Darmstadt, Germany).

**Amplicon Sequencing**

The final DNA pool spiked with 10% PhiX genomic DNA (Illumina, San Diego, CA) was used for MiSeq sequencing (Illumina, San Diego, CA) using the MiSeq Reagent Kit v3 (Illumina, San Diego, CA). 0.5 µM of forward, reverse and index sequencing primers complementary to the linker/primer region of the concatenated primers were added together (sequence of primers are available in Suppl Table 3). Paired-end sequencing was performed for 600 cycles for 301 bp amplicon size according to the Illumina instructions (http://os.bioprotocol.org/attached/file/20171217/miseq%20denature%20dilute%20libraries%20guide%2015039740%2003.pdf).

Table S1

| Year | Location | Species | Latitude | Longitude |
|---|---|---|---|---|
| 2017 | Turkey,Kışlak | Ta (non-treated) | 35° 58' 21.8'' | 36° 07' 41.8'' |
| 2017 | Germany | Ta (benchmark) | 54°18'50.9" | 9°59'42.2" |
| 2005 | West population | Tb | 37° 19' 31'' | 37° 09' 28'' |
| 2006 | West population | Tb | 36°54'29" | 37°11'49" |
| 2006 | West population | Td | 37°20'19" | 37°16'50" |
| 2006 | West population | Td | 37°19'50" | 37°18'51" |
| 2004 | East population | Td | 37°50'40" | 39°47'58" |
| 2004 | East population | Tu | 37°50'40" | 39°47'58" |
| 2004 | East population | Td | 37°47'31" | 39°57'18" |

Table S2

| | # of samples | Total # of reads | Average # of reads | # of OTUs (b.r.) | # of OTUs (a.r.) |
|---|---|---|---|---|---|
| **Seeds- bacteria** | 58 | 2356761 | 40633.81 | 1441 | 1157 |
| **Seeds-fungi** | 30 | 2621026 | 87367.53 | 272 | 124 |
| **Axenic leaves- bacteria** | 16 | 633975 | 84526.0 | 760 | 589 |
| **Axenic leaves- fungi** | 15 | 719618 | 47974.53 | 119 | 98 |
| **Axenic roots- bacteria** | 16 | 663813 | 41488.31 | 828 | 632 |
| **Axenic roots- fungi** | 16 | 2309064 | 144316.5 | 113 | 74 |
| **Leaves- bacteria** | 42 | 2460327 | 58579.21 | 3864 | 3676 |
| **Roots- bacteria** | 39 | 1690934 | 43357.28 | 3184 | 3100 |
| **Roots-fungi** | 42 | 1545621 | 36800.5 | 3757 | 3737 |
| **Natural soil- bacteria** | 3 | 100190 | 33396.67 | 1258 | 1248 |
| **Agricultural soil-bacteria** | 3 | 82333 | 27444.33 | 1270 | 1266 |

Table S3

16S, ITS universal and interfering primers

| Universal_Primers- PCR1 |
|---|
| Bacteria - V5/V6/V7 |
| 799F: AACMGGATTAGATACCCKG |
| 1192R: ACGTCATCCCCACCTTCC |
| Fungi ITS 1- PCR1 |
| ITS1F: CTTGGTCATTTAGAGGAAGTAA |
| ITS2: GCTGCGTTCTTCATCGATGC |
| Interfering Primers: Mitochondria - V5/V6/V7 |
| Forward Primer: GGATCAGGGGCCCAGCTAACGCGTGAAACA |
| Reverse Primer: CGGAGCGGGGCGCGTACTATTACCACTACG |

Bacteria 16S, ITS PCR2 primers

| |
|---|
| B5-F: AATGATACGGCGACCACCGAGATCTACACGACTGCGACTGGCGAACMGGATTAGATACCCKG |
| B5-R: CAAGCAGAAGACGGCATACGAGATXXXXXXXXXXXXCAGCCATTTAGTGTCACGTCATCCCCACCTTCC |
| Fungi, XXXXXXXXXXXX: 12bp barcode sequence |
| ITS-F: AATGATACGGCGACCACCGAGATCTACACTCACGCGCAGGCTTGGTCATTTAGAGGAAGTAA |
| ITS-R: CAAGCAGAAGACGGCATACGAGATXXXXXXXXXXXXCGTACTGTGGAGAGCTGCGTTCTTCATCGATGC |

16S, ITS sequencing primers

| |
|---|
| B5-R1 |
| ACGACTGCGACTGGCGAACMGGATTAGATACCC |
| B5-R2 |
| CAGCCATTTAGTGTCACGTCATCCCCACCTTCC |

| |
|---|
| B5-R3 |
| GGAAGGTGGGGATGACGTGACACTAAATGGCTG |
| ITS-R1 |
| TCACGCGCAGGCTTGGTCATTTAGAGGAAGTAA |
| ITS-R2 |
| CGTACTGTGGAGAGCTGCGTTCTTCATCGATGC |
| ITS-R3 |
| GCATCGATGAAGAACGCAGCTCTCCACAGTACG |

Suppl Fig 1

A

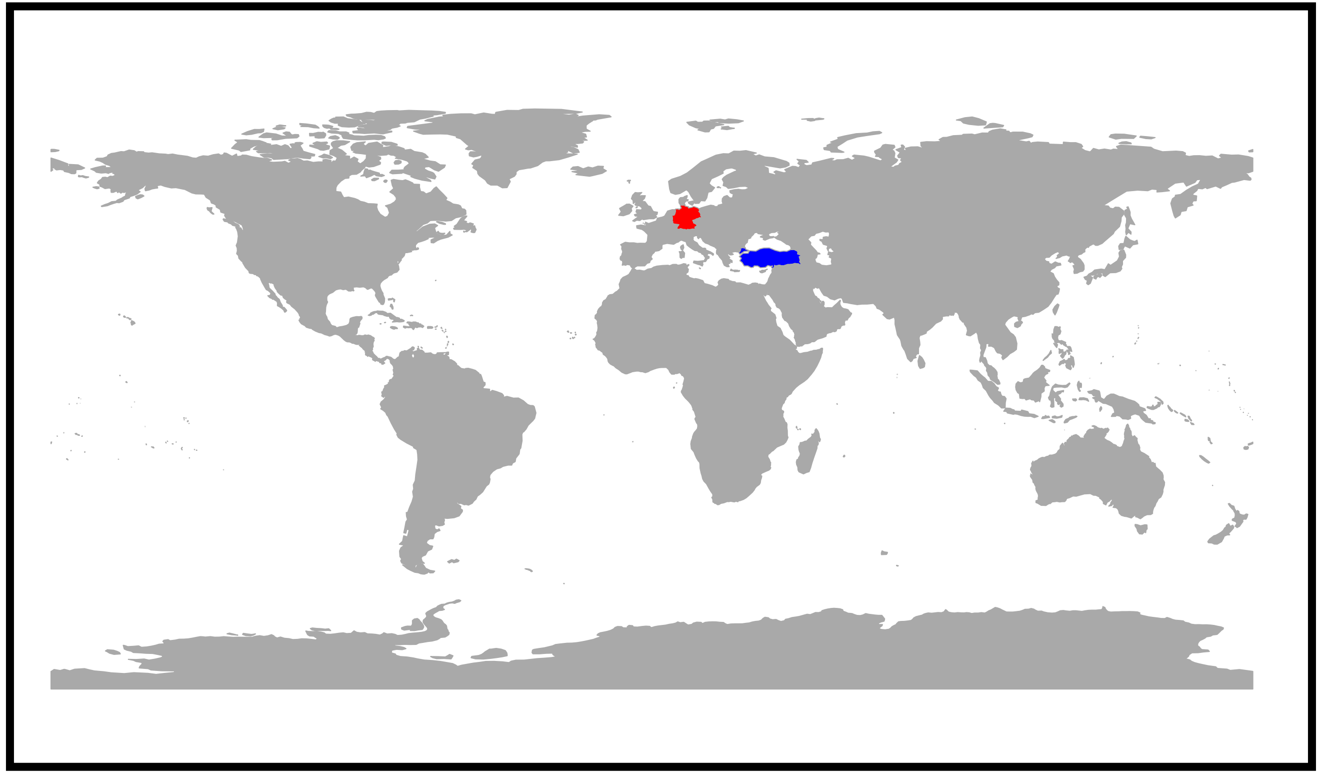

B

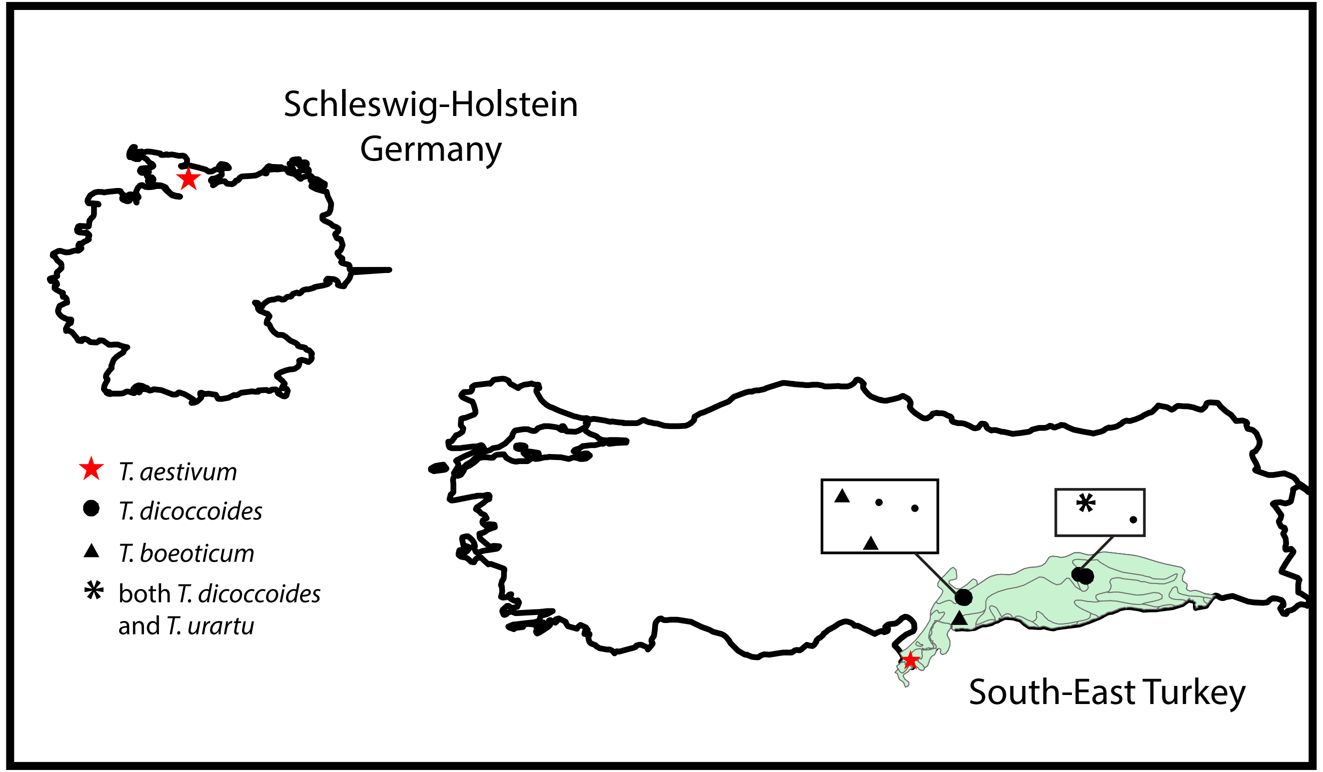

Suppl Fig 2

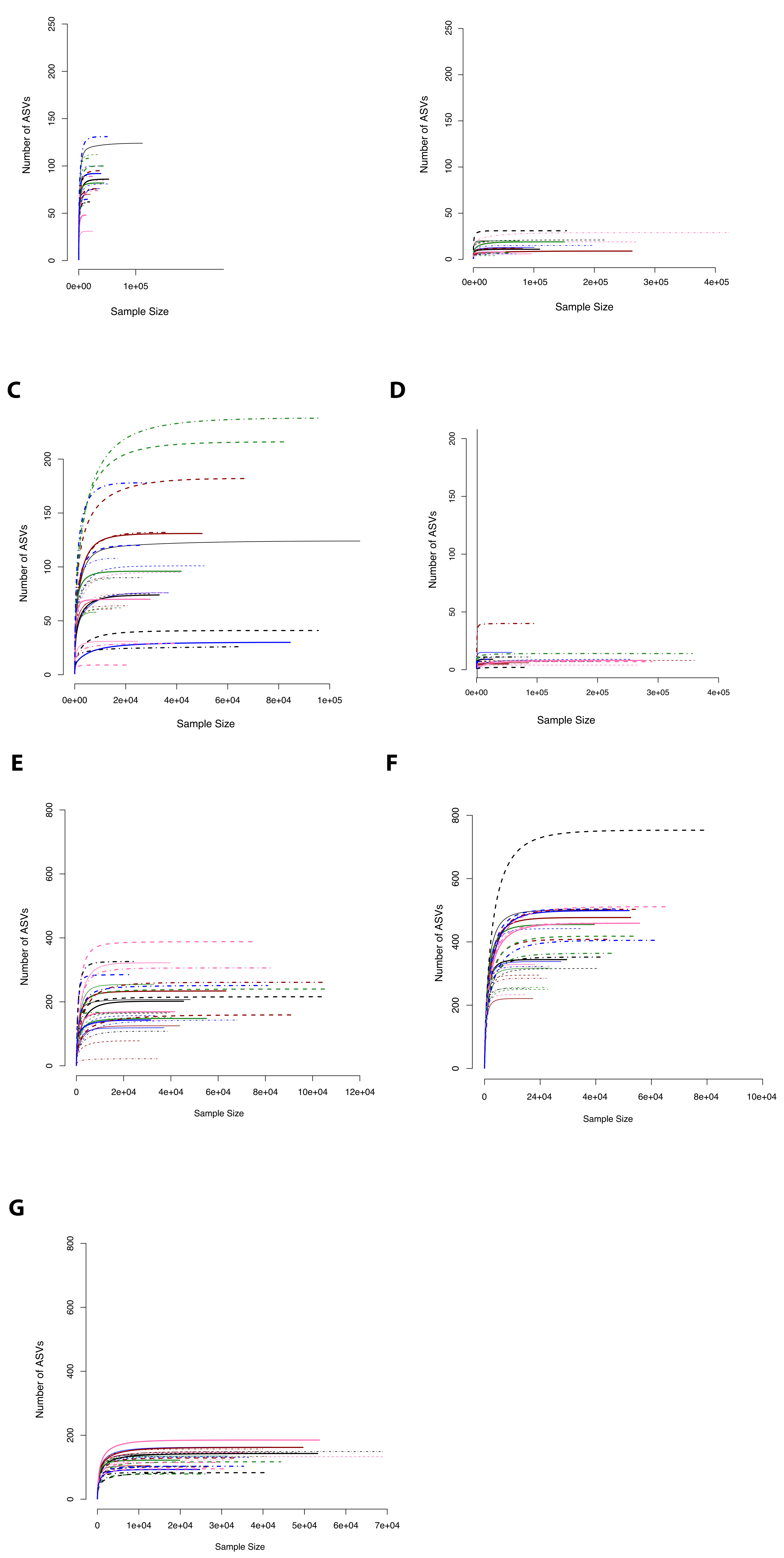
A
Number of ASVs
0
50
100
150
200
250
0e+00
1e+05
Sample Size
B
Number of ASVs
0
50
100
150
200
250
0e+00
1e+05
2e+05
3e+05
4e+05
Sample Size
C
Number of ASVs
0
50
100
150
200
0e+00
2e+04
4e+04
6e+04
8e+04
1e+05
Sample Size
D
Number of ASVs
0
50
100
150
200
0e+00
1e+05
2e+05
3e+05
4e+05
Sample Size
E
Number of ASVs
0
200
400
600
800
0
2e+04
4e+04
6e+04
8e+04
10e+04
12e+04
Sample Size
F
Number of ASVs
0
200
400
600
800
0
24+04
4e+04
6e+04
8e+04
10e+04
Sample Size
G
Number of ASVs
0
200
400
600
800
0
1e+04
2e+04
3e+04
4e+04
5e+04
6e+04
7e+04
Sample Size

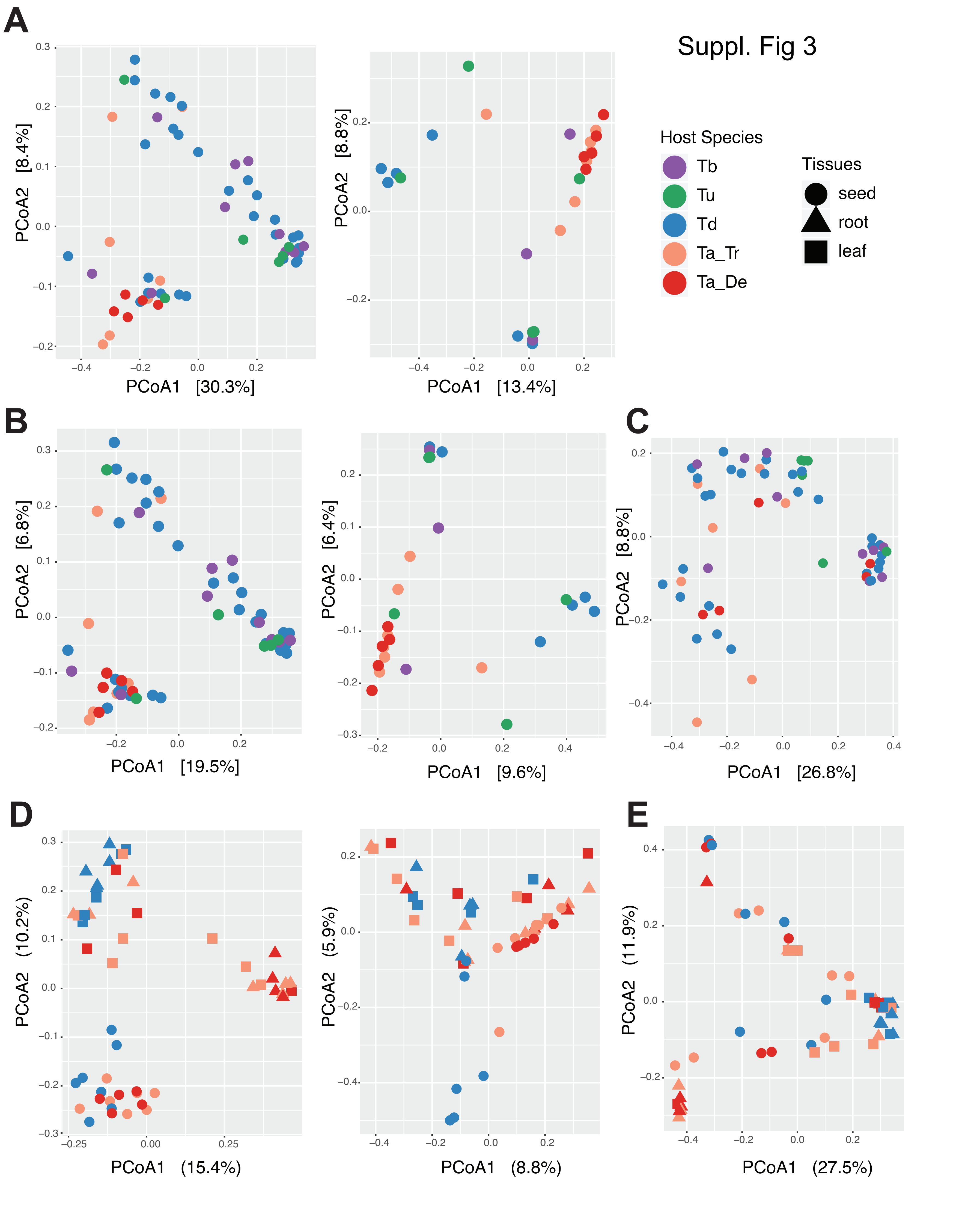

Suppl. Fig 3
A
B
C
D
E
Host Species
Tb
Tu
Td
Ta_Tr
Ta_De
Tissues
seed
root
leaf
PCoA1 [30.3%]
PCoA2 [8.4%]
PCoA1 [13.4%]
PCoA2 [8.8%]
PCoA1 [19.5%]
PCoA2 [6.8%]
PCoA1 [9.6%]
PCoA2 [6.4%]
PCoA1 [26.8%]
PCoA2 [8.8%]
PCoA1 (15.4%)
PCoA2 (10.2%)
PCoA1 (8.8%)
PCoA2 (5.9%)
PCoA1 (27.5%)
PCoA2 (11.9%)

Suppl Fig 4

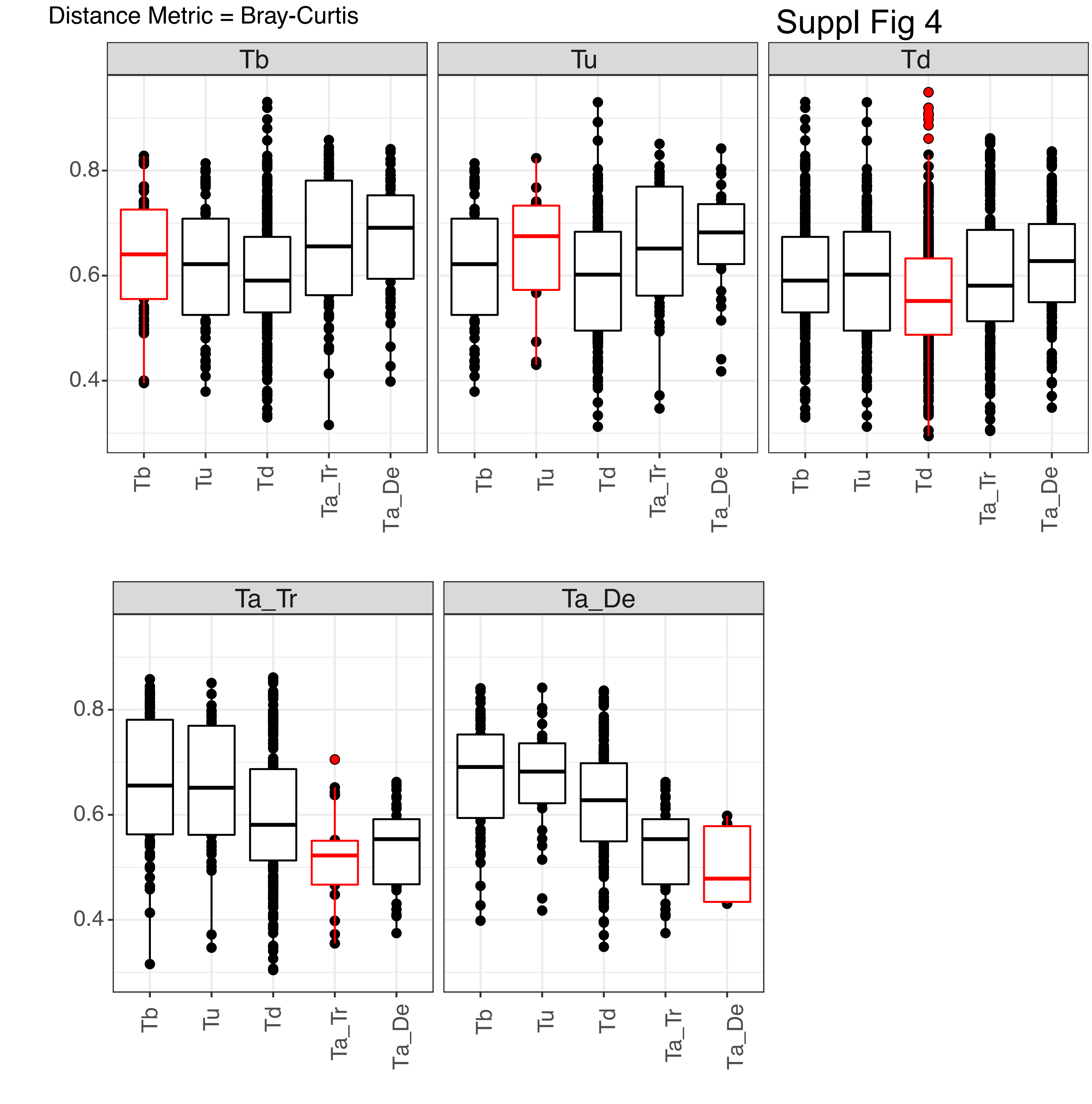
Distance Metric = Bray-Curtis
Tb
Tu
Td
Ta_Tr
Ta_De
0.4
0.6
0.8
Tb
Tu
Td
Ta_Tr
Ta_De

Distance Metric = Unweighted UniFrac

Suppl Fig 5

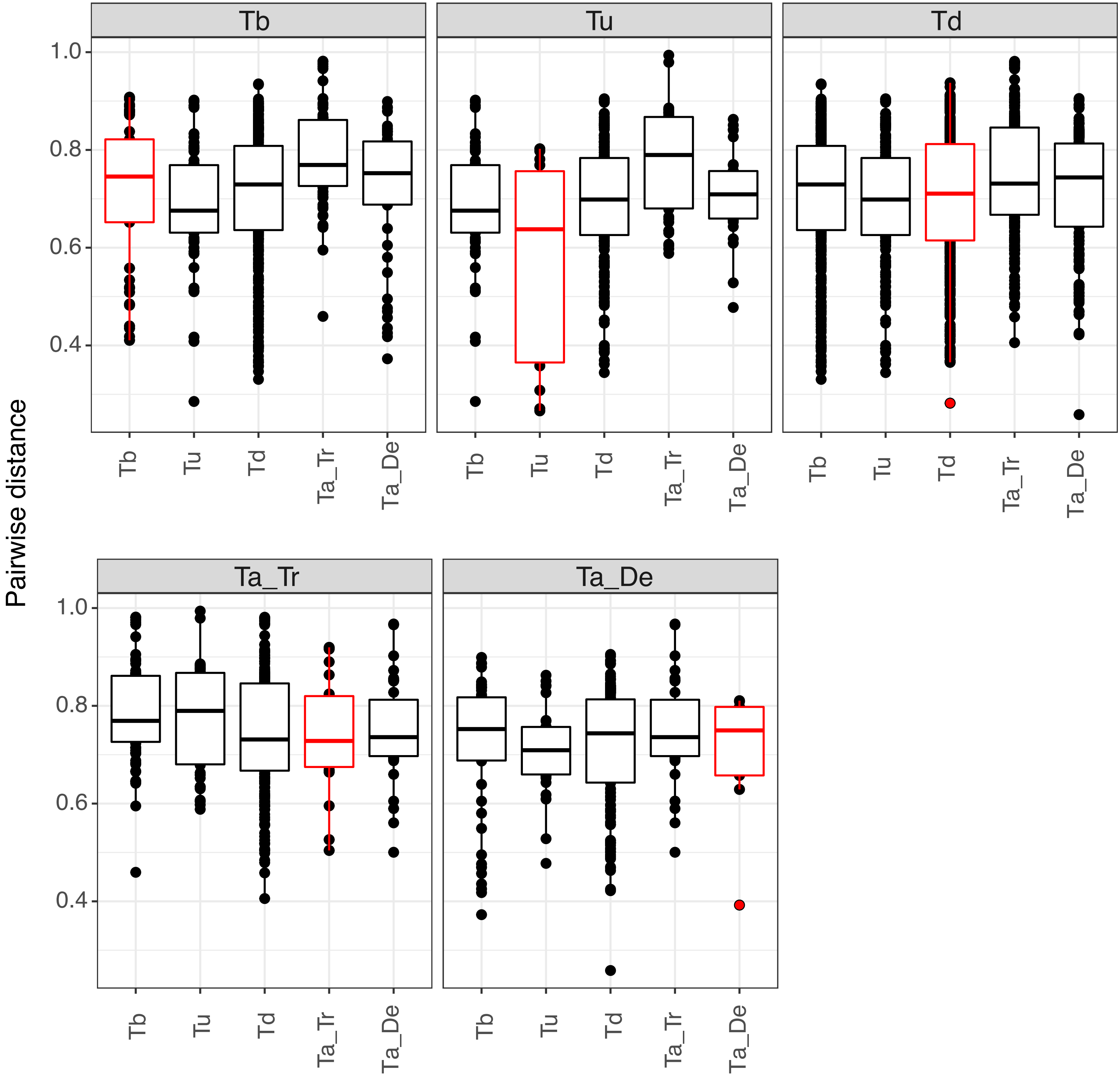

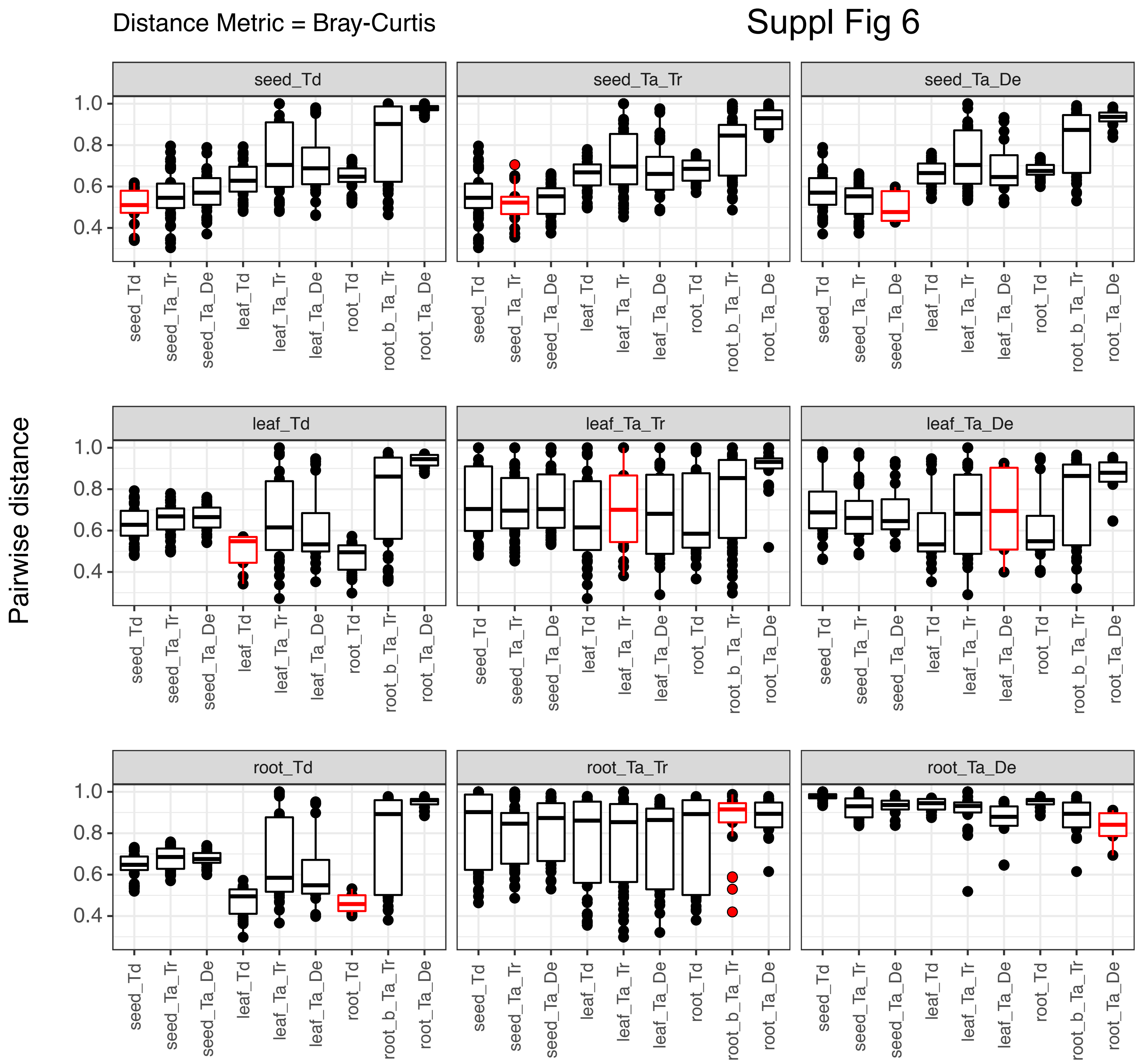

Distance Metric = Bray-Curtis
Suppl Fig 6
Pairwise distance
seed_Td
seed_Ta_Tr
seed_Ta_De
leaf_Td
leaf_Ta_Tr
leaf_Ta_De
root_Td
root_Ta_Tr
root_Ta_De
root_b_Ta_Tr
0.4
0.6
0.8
1.0

### Suppl Fig 7

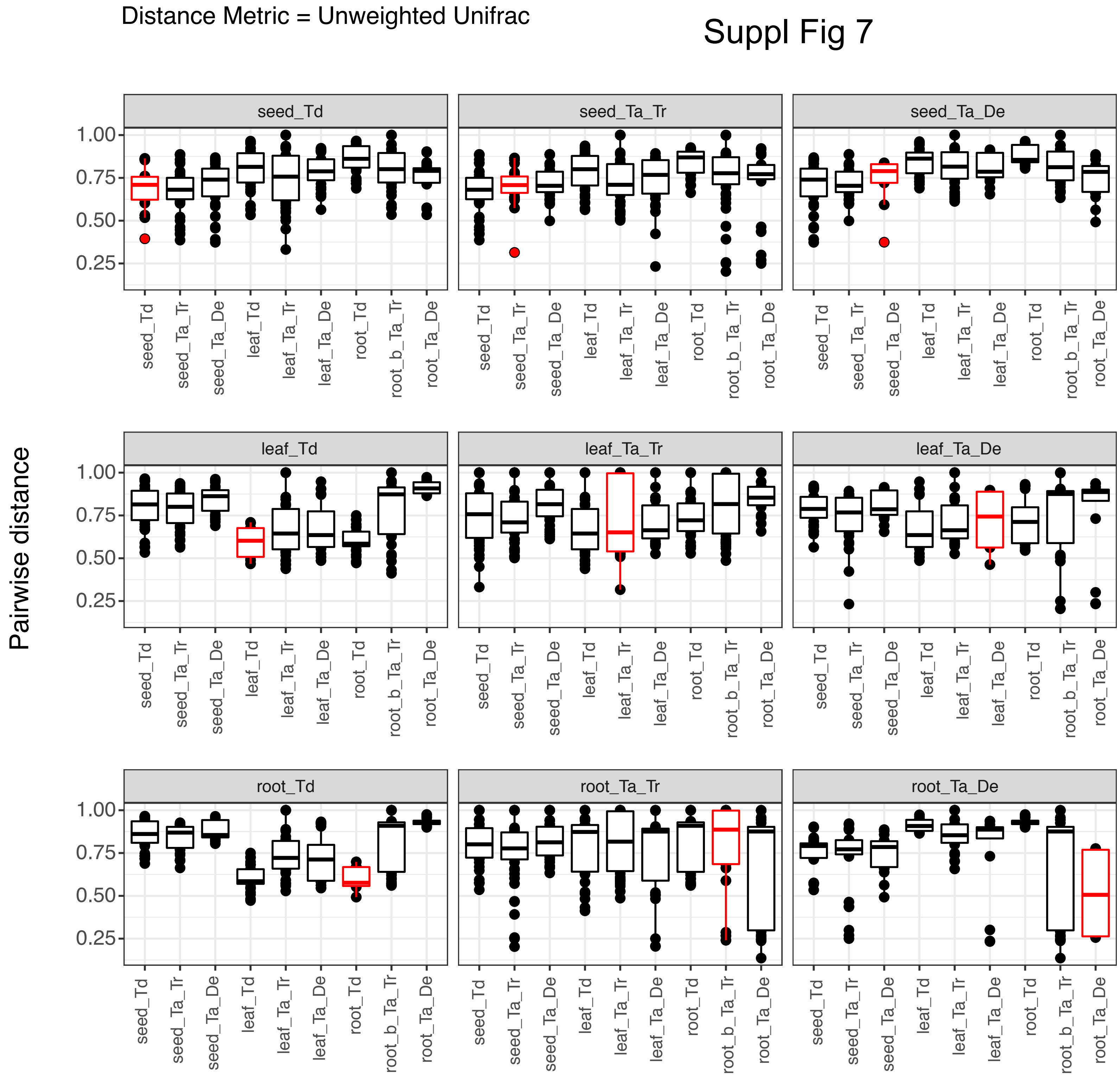
Distance Metric = Unweighted Unifrac
Pairwise distance
seed_Td
seed_Ta_Tr
seed_Ta_De
leaf_Td
leaf_Ta_Tr
leaf_Ta_De
root_Td
root_Ta_Tr
root_Ta_De
1.00
0.75
0.50
0.25
root_b_Ta_Tr

Suppl Fig 8

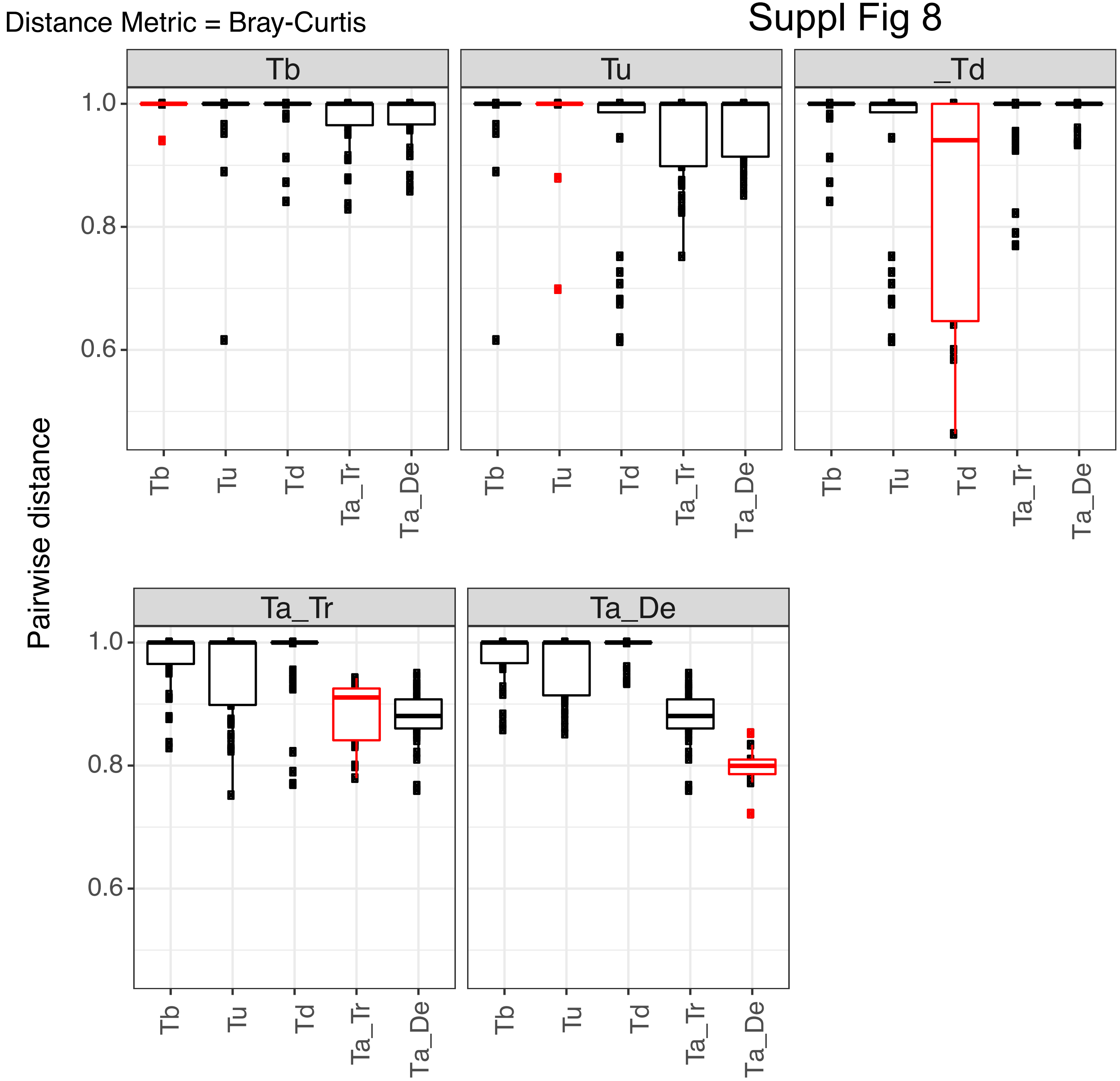

Distance Metric = Bray-Curtis
Pairwise distance
Tb
Tu
_Td
Ta_Tr
Ta_De
1.0
0.8
0.6
Tb
Tu
Td
Ta_Tr
Ta_De

Suppl Fig 9

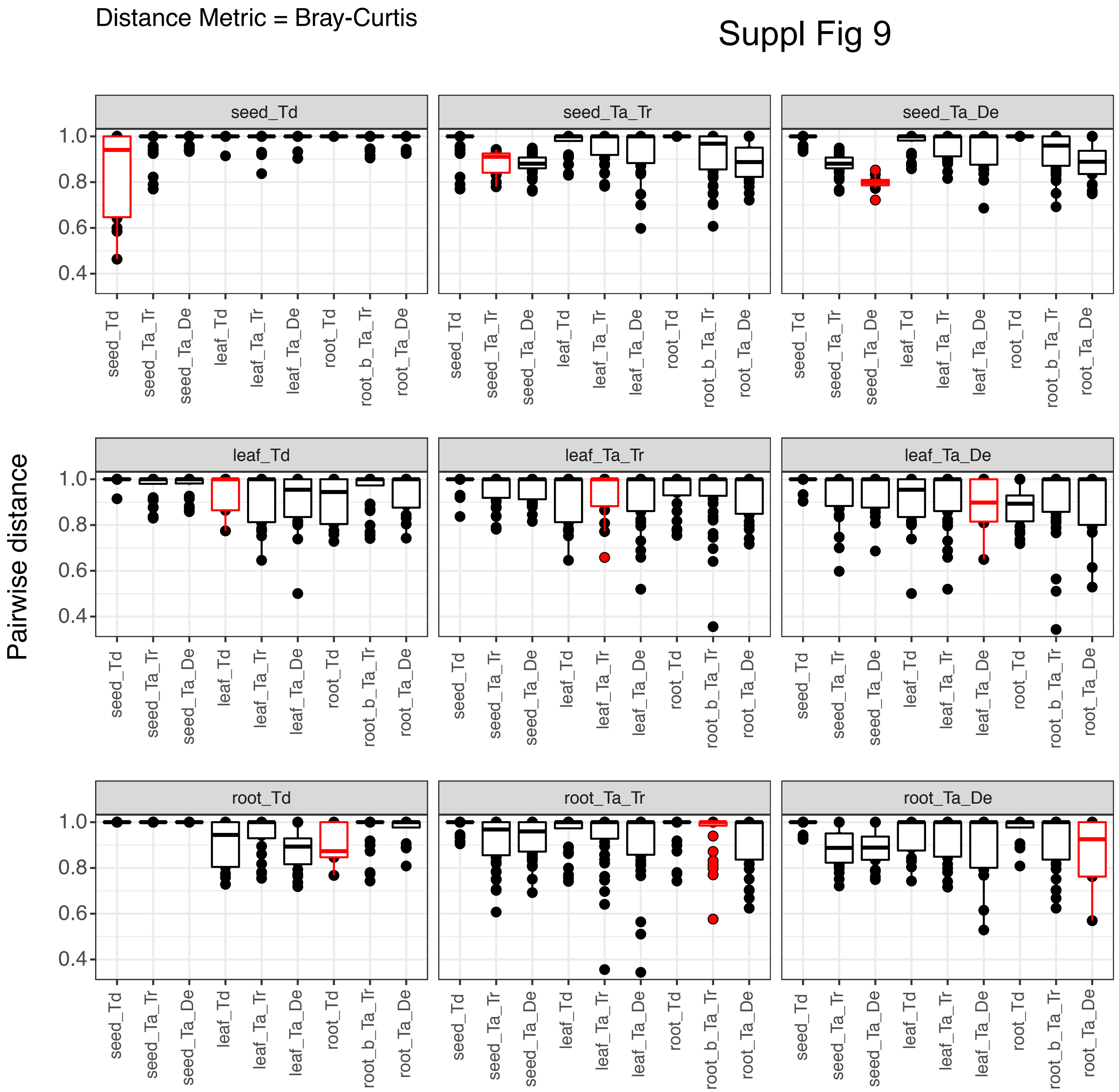

Distance Metric = Bray-Curtis
Pairwise distance
seed_Td
seed_Ta_Tr
seed_Ta_De
leaf_Td
leaf_Ta_Tr
leaf_Ta_De
root_Td
root_Ta_Tr
root_Ta_De
1.0
0.8
0.6
0.4
seed_Td
seed_Ta_Tr
seed_Ta_De
leaf_Td
leaf_Ta_Tr
leaf_Ta_De
root_Td
root_b_Ta_Tr
root_Ta_De

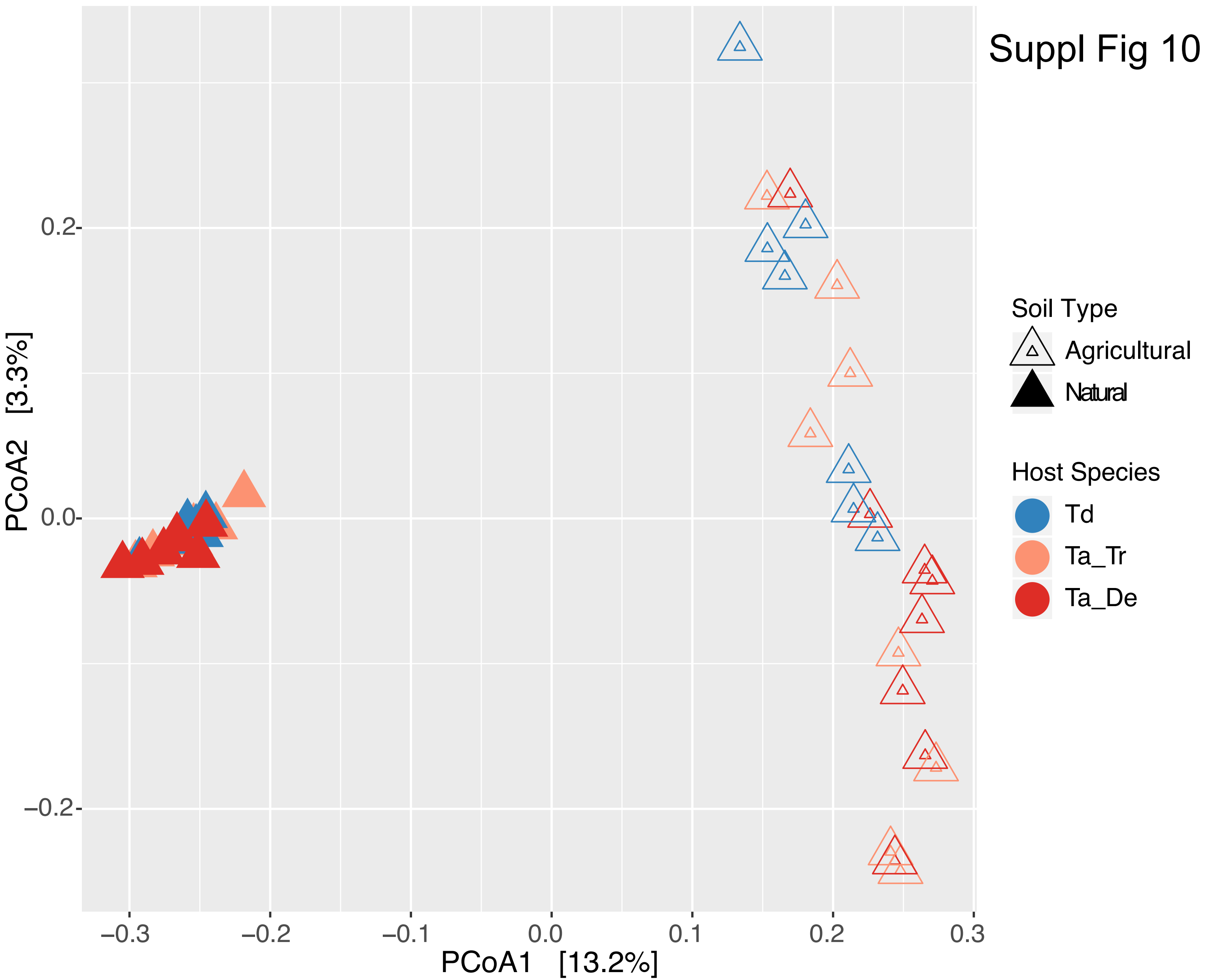

Suppl Fig 10
PCoA2 [3.3%]
0.2
0.0
−0.2
−0.3
−0.2
−0.1
0.0
0.1
0.2
0.3
PCoA1 [13.2%]
Soil Type
Agricultural
Natural
Host Species
Td
Ta_Tr
Ta_De

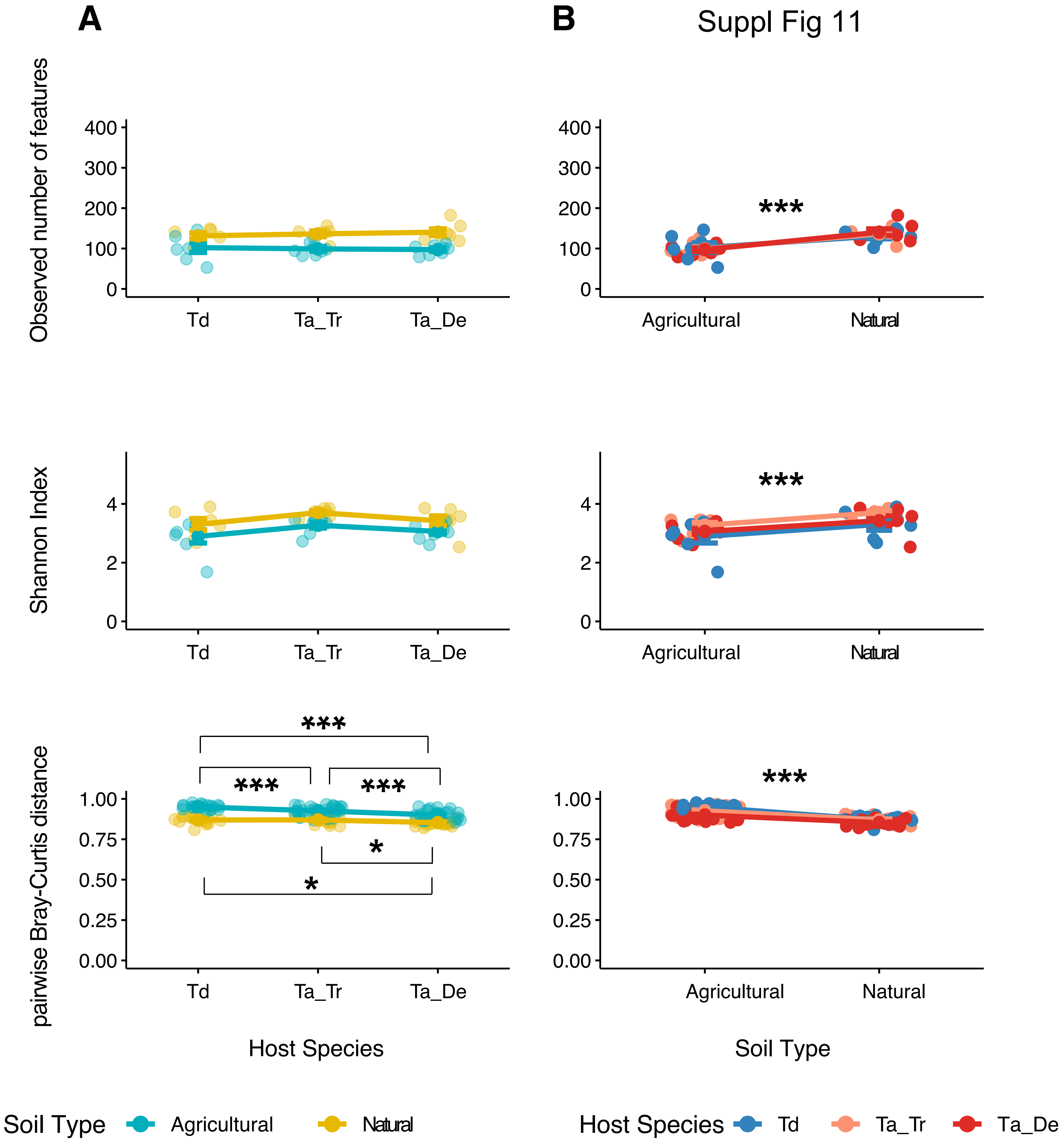
A
B
Suppl Fig 11
Observed number of features
Shannon Index
pairwise Bray-Curtis distance
400
300
200
100
0
4
2
0
1.00
0.75
0.50
0.25
0.00
Td
Ta_Tr
Ta_De
Agricultural
Natural
***
*
Host Species
Soil Type
Soil Type
Agricultural
Natural
Host Species
Td
Ta_Tr
Ta_De

Suppl Fig 12

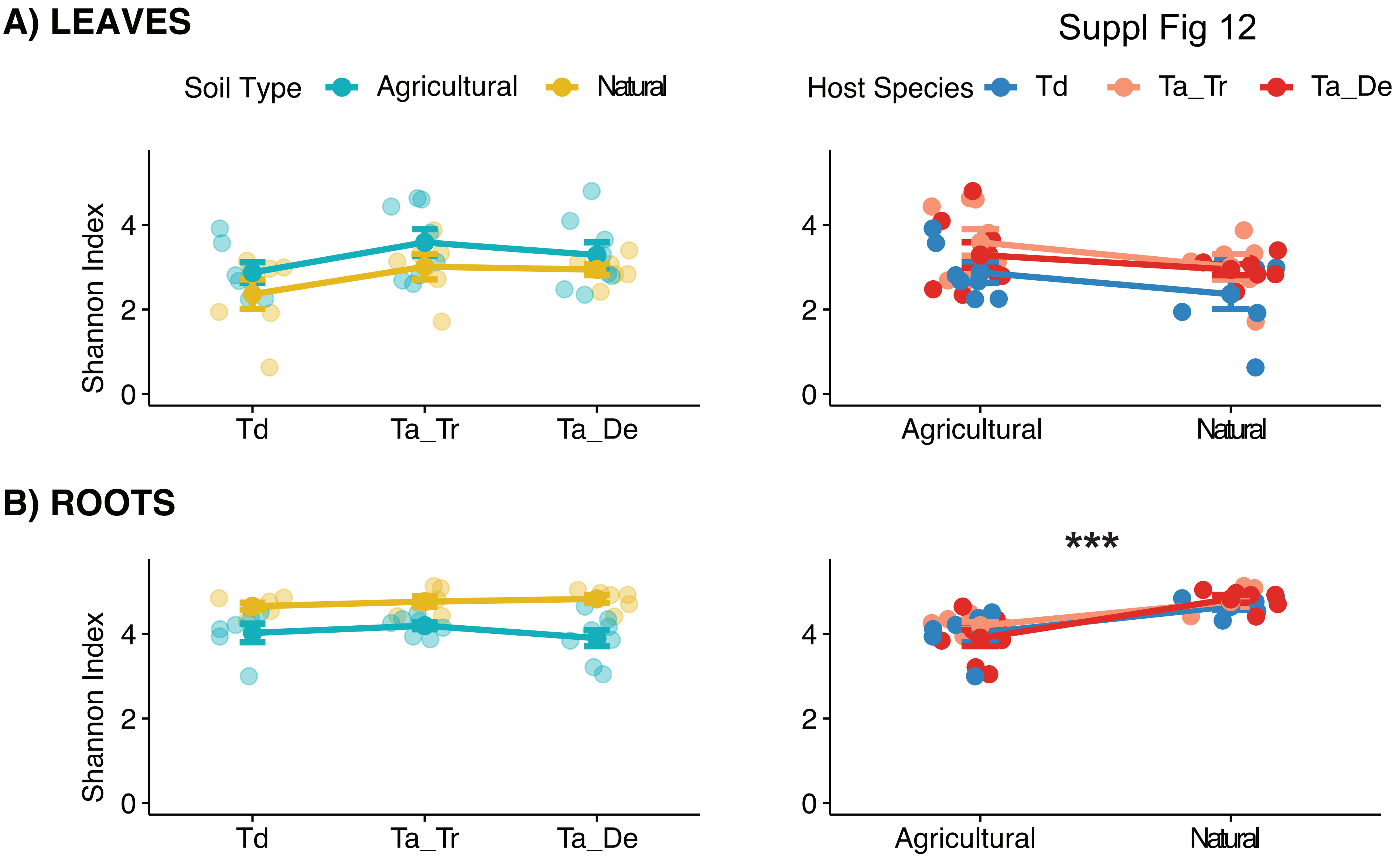

A) LEAVES
Soil Type
Agricultural
Natural
Host Species
Td
Ta_Tr
Ta_De
Shannon Index
0
2
4
Td
Ta_Tr
Ta_De
Agricultural
Natural
B) ROOTS
***

Suppl Fig 13

## A

| | Leaf Td | Leaf Ta_Tr | Leaf Ta_De | Root Td | Root Ta_Tr | Root Ta_De | Soil natural soil | Soil agr. soil |
|---|---|---|---|---|---|---|---|---|
| Proteobacteria; Oxalobacteraceae | 22.2 | 12.2 | 21.1 | 18.6 | 26.5 | 24.4 | 1 | 0.9 |
| Actinobacteria; Streptomycetaceae | 0.3 | 2 | 3 | 32.6 | 27 | 30 | 0.7 | 10.3 |
| Proteobacteria; Comamonadaceae | 7.4 | 19.9 | 16.1 | 8.3 | 9.8 | 8.8 | 2 | 3.5 |
| Proteobacteria; Rhizobiaceae | 21.3 | 12.8 | 12.1 | 1 | 1.3 | 1.1 | 0.2 | 0 |
| Proteobacteria; Halomonadaceae | 13.6 | 15.9 | 15.7 | 0 | 0.1 | 0 | 0 | 0 |
| Proteobacteria; Vibrionaceae | 14.7 | 14 | 8.4 | 0 | 0.1 | 0 | 0 | 0 |
| Actinobacteria; Micromonosporaceae | 0.1 | 0.5 | 0.6 | 5.6 | 5.9 | 7.7 | 3.4 | 1.9 |
| Actinobacteria; Nocardioidaceae | 0.3 | 0.9 | 1.5 | 5.1 | 4.2 | 4 | 4 | 5.4 |
| Bacteroidetes; Flavobacteriaceae | 0.2 | 0.4 | 0.2 | 5.9 | 4.6 | 4.7 | 0.3 | 2.1 |
| Proteobacteria; Xanthomonadaceae | 1.3 | 1.6 | 1.3 | 3 | 2.7 | 2.7 | 1.5 | 5.9 |
| Proteobacteria; Burkholderiaceae | 3.5 | 2 | 3.2 | 0.4 | 0.5 | 0.3 | 0.1 | 2.1 |
| Proteobacteria; Phyllobacteriaceae | 3.5 | 1.6 | 3.1 | 0.3 | 0.3 | 0.4 | 0.2 | 0.5 |
| Proteobacteria; Pseudomonadaceae | 2.7 | 1.7 | 1.6 | 0.6 | 0.6 | 0.5 | 0.2 | 0.9 |
| Actinobacteria; Geodermatophilaceae | 0 | 0.2 | 0.2 | 0.5 | 0.5 | 0.4 | 21.8 | 0.2 |
| Proteobacteria; Hyphomicrobiaceae | 0.2 | 0.4 | 0.3 | 1.4 | 1.4 | 1.4 | 0.7 | 6.9 |
| Actinobacteria; Actinosynnemataceae | 0 | 0.1 | 0.1 | 2.5 | 1.8 | 1.9 | 1.6 | 0.4 |
| Proteobacteria; Caulobacteraceae | 0.1 | 0.2 | 0.2 | 1.7 | 1.6 | 1.4 | 1 | 1.6 |
| Actinobacteria; Rubrobacteraceae | 0 | 0.2 | 0.1 | 0.3 | 0.2 | 0.2 | 12.9 | 0.1 |
| Actinobacteria; Microbacteriaceae | 0.1 | 0.2 | 0.3 | 1 | 0.8 | 0.9 | 1.4 | 1.5 |
| Proteobacteria; [Chromatiaceae] | 1.4 | 1.5 | 0.6 | 0 | 0.1 | 0 | 0 | 0 |

#### B LEAVES

| | Agricultural Td | Agricultural Ta_Tr | Agricultural Ta_De | Natural Td | Natural Ta_Tr | Natural Ta_De |
|---|---|---|---|---|---|---|
| Proteobacteria; Oxalobacteraceae | 44.1 | 20.3 | 36.2 | 0.3 | 1.3 | 1 |
| Proteobacteria; Rhizobiaceae | 0.1 | 0.1 | 0 | 42.5 | 29.6 | 28.2 |
| Proteobacteria; Halomonadaceae | 11.2 | 10.5 | 8.1 | 16.1 | 23.2 | 25.8 |
| Proteobacteria; Comamonadaceae | 10.4 | 21.3 | 17.1 | 4.4 | 18.1 | 14.8 |
| Proteobacteria; Vibrionaceae | 8.1 | 16.3 | 4.5 | 21.2 | 11 | 13.6 |
| Proteobacteria; Burkholderiaceae | 7 | 2.9 | 5.6 | 0 | 0.7 | 0.1 |
| Proteobacteria; Phyllobacteriaceae | 0.1 | 0.1 | 0.2 | 7 | 3.7 | 6.9 |
| Proteobacteria; Pseudomonadaceae | 4.4 | 2.6 | 2.4 | 1.1 | 0.4 | 0.4 |
| Actinobacteria; Streptomycetaceae | 0.5 | 3.1 | 5.1 | 0 | 0.6 | 0.1 |
| Proteobacteria; Xanthomonadaceae | 1.5 | 2.2 | 1.9 | 1 | 0.8 | 0.6 |
| Proteobacteria; [Chromatiaceae] | 0.6 | 2.1 | 0.2 | 2.3 | 0.8 | 1.3 |
| Proteobacteria; Neisseriaceae | 1.5 | 1.7 | 1.8 | 0.7 | 0.5 | 0.7 |
| Actinobacteria; Nocardioidaceae | 0.5 | 1.2 | 2.4 | 0 | 0.6 | 0.3 |
| Actinobacteria; Propionibacteriaceae | 0.8 | 2.1 | 0.8 | 0.1 | 0.3 | 0.3 |
| Firmicutes; Streptococcaceae | 0.8 | 1.1 | 0.7 | 0.6 | 0.3 | 0.5 |
| Proteobacteria; Enterobacteriaceae | 0.5 | 1.1 | 0.5 | 0.7 | 0.4 | 0.5 |
| Proteobacteria; Methylobacteriaceae | 0.5 | 0.9 | 0.5 | 0.1 | 0.7 | 0.5 |
| Actinobacteria; Micromonosporaceae | 0.1 | 0.5 | 1 | 0.1 | 0.5 | 0.1 |
| Firmicutes; Staphylococcaceae | 0.3 | 0.7 | 0.7 | 0 | 0.2 | 0.2 |
| Proteobacteria; Methylophilaceae | 0.2 | 0.5 | 0.6 | 0.1 | 0.4 | 0.2 |

#### C ROOTS

| | Agricultural Td | Agricultural Ta_Tr | Agricultural Ta_De | Natural Td | Natural Ta_Tr | Natural Ta_De |
|---|---|---|---|---|---|---|
| Actinobacteria; Streptomycetaceae | 49.7 | 40.2 | 46.2 | 15.5 | 11.5 | 8.5 |
| Proteobacteria; Oxalobacteraceae | 11.6 | 22 | 23.3 | 25.7 | 31.8 | 25.9 |
| Proteobacteria; Comamonadaceae | 2.7 | 5.2 | 3.3 | 14 | 15 | 16.1 |
| Actinobacteria; Micromonosporaceae | 3.6 | 3.5 | 3.3 | 7.6 | 8.8 | 13.6 |
| Bacteroidetes; Flavobacteriaceae | 0.5 | 0.5 | 0.3 | 11.2 | 9.4 | 10.5 |
| Actinobacteria; Nocardioidaceae | 8 | 6.2 | 5.4 | 2.2 | 1.8 | 2.1 |
| Proteobacteria; Xanthomonadaceae | 4.1 | 3.6 | 3.1 | 2 | 1.6 | 2.2 |
| Actinobacteria; Actinosynnemataceae | 1.4 | 0.9 | 0.5 | 3.7 | 2.9 | 3.9 |
| Proteobacteria; Caulobacteraceae | 1.7 | 1.5 | 1 | 1.7 | 1.7 | 1.8 |
| Proteobacteria; Hyphomicrobiaceae | 2 | 2 | 1.7 | 0.7 | 0.7 | 0.9 |
| Proteobacteria; Rhizobiaceae | 0.6 | 1.2 | 0.8 | 1.5 | 1.4 | 1.5 |
| Actinobacteria; Thermomonosporaceae | 1 | 1.2 | 1.1 | 1.1 | 1.3 | 0.8 |
| Actinobacteria; Microbacteriaceae | 1 | 0.8 | 0.7 | 0.9 | 0.8 | 1.2 |
| Actinobacteria; Promicromonosporaceae | 0.4 | 0.5 | 0.3 | 1.7 | 0.9 | 1.4 |
| Chloroflexi; [Kouleothrixaceae] | 0.4 | 0.4 | 0.4 | 0.7 | 1.4 | 1.1 |
| Actinobacteria; Pseudonocardiaceae | 0.3 | 0.4 | 0.1 | 1.1 | 1.1 | 0.9 |
| Proteobacteria; Pseudomonadaceae | 0.5 | 0.4 | 0.2 | 0.8 | 0.9 | 0.8 |
| Firmicutes; Paenibacillaceae | 0.8 | 1.3 | 0.8 | 0 | 0 | 0 |
| Proteobacteria; Bradyrhizobiaceae | 0.8 | 0.7 | 0.5 | 0.3 | 0.4 | 0.4 |
| Actinobacteria; Micrococcaceae | 0.5 | 0.2 | 0.5 | 1 | 0.5 | 0.4 |

high — low

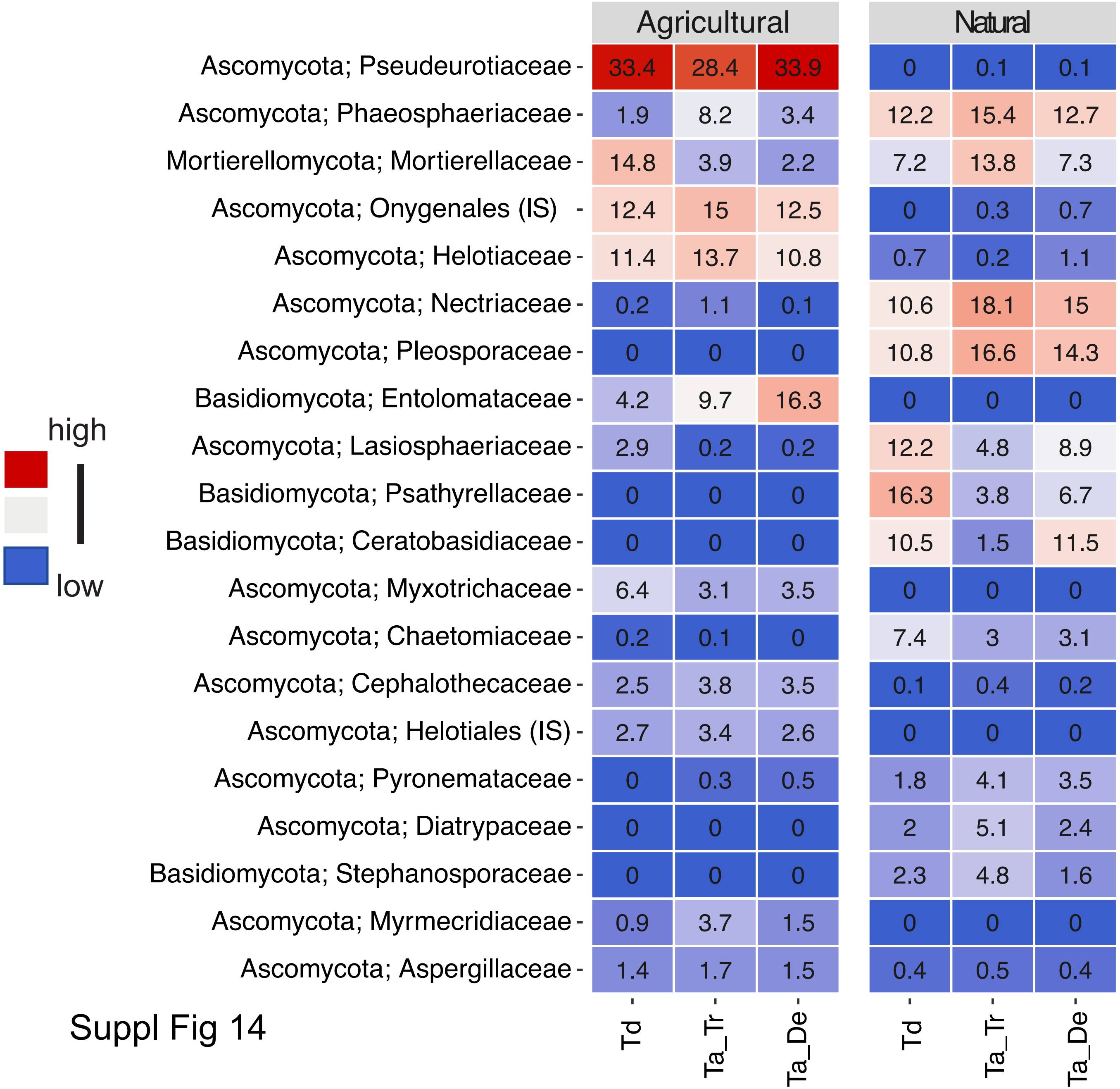
Agricultural
Natural
Ascomycota; Pseudeurotiaceae 33.4 28.4 33.9 0 0.1 0.1
Ascomycota; Phaeosphaeriaceae 1.9 8.2 3.4 12.2 15.4 12.7
Mortierellomycota; Mortierellaceae 14.8 3.9 2.2 7.2 13.8 7.3
Ascomycota; Onygenales (IS) 12.4 15 12.5 0 0.3 0.7
Ascomycota; Helotiaceae 11.4 13.7 10.8 0.7 0.2 1.1
Ascomycota; Nectriaceae 0.2 1.1 0.1 10.6 18.1 15
Ascomycota; Pleosporaceae 0 0 0 10.8 16.6 14.3
Basidiomycota; Entolomataceae 4.2 9.7 16.3 0 0 0
Ascomycota; Lasiosphaeriaceae 2.9 0.2 0.2 12.2 4.8 8.9
Basidiomycota; Psathyrellaceae 0 0 0 16.3 3.8 6.7
Basidiomycota; Ceratobasidiaceae 0 0 0 10.5 1.5 11.5
Ascomycota; Myxotrichaceae 6.4 3.1 3.5 0 0 0
Ascomycota; Chaetomiaceae 0.2 0.1 0 7.4 3 3.1
Ascomycota; Cephalothecaceae 2.5 3.8 3.5 0.1 0.4 0.2
Ascomycota; Helotiales (IS) 2.7 3.4 2.6 0 0 0
Ascomycota; Pyronemataceae 0 0.3 0.5 1.8 4.1 3.5
Ascomycota; Diatrypaceae 0 0 0 2 5.1 2.4
Basidiomycota; Stephanosporaceae 0 0 0 2.3 4.8 1.6
Ascomycota; Myrmecridiaceae 0.9 3.7 1.5 0 0 0
Ascomycota; Aspergillaceae 1.4 1.7 1.5 0.4 0.5 0.4
Td
Ta_Tr
Ta_De
Td
Ta_Tr
Ta_De
high
low

Suppl Fig 14
